# Striatal Cholinergic Interneurons Signal Aversion and Drive Avoidance

**DOI:** 10.64898/2026.05.27.728172

**Authors:** Yuxin Zhou, Hua-an Tseng, Jiancheng Xie, Erynne San Antonio, Aiden Ly, Abdullah Malik, Christopher Fiorillo, Xue Han

**Affiliations:** Department of Biomedical Engineering, Boston University, Boston, MA, USA; Department of Bio and Brain Engineering, Korea Advanced Institute of Science and Technology (KAIST), Daejeon, Republic of Korea

**Keywords:** cholinergic interneurons, voltage imaging, ElectraOff, SomArchon, ArchT, aversive and reward, caudate and putamen

## Abstract

Seeking reward and avoiding punishment are essential for survival. However, how the brain learns aversive value and generates avoidance remains poorly understood. We investigate acetylcholine-releasing striatal cholinergic interneurons (ChIs), which exhibit prominent excitation to salient behavioral cues but have largely been postulated to play a permissive role in reinforcement learning. Using voltage imaging in mice performing aversive versus reward conditioning tasks, we reveal learning-mediated changes in both subthreshold membrane voltage and suprathreshold spiking of individual ChIs. Across tasks, ChIs exhibit stronger immediate excitation to conditioned cues predicting negative outcomes than positive outcomes. This valence-dependent response augmentation reflects a learning-induced network change specific to cues predicting negative outcomes. Furthermore, brief optogenetic silencing of this immediate excitation selectively impairs conditioned avoidance, but not conditioned approach. Together, these results reveal that ChIs actively signal evidence for aversion and play a critical role in driving avoidance, highlighting a cholinergic mechanism underlying aversive learning.

## Introduction

Approach and avoidance are initiated based on the value of sensory cues. Value learning engages broad sensorimotor areas to optimize reward and minimize punishment. However, how the brain learns aversion remains poorly understood^1,2^. The dorsal striatum is critical for sensorimotor decision making and has high concentrations of two key neuromodulators, acetylcholine (ACh) released by local cholinergic interneurons (ChIs) and dopamine by midbrain dopaminergic neurons^3,4^. While there is a well-developed framework of dopamine in signaling reward prediction errors and facilitating approach behavior^5-9^, the role of dopamine suppression in avoidance remains controversial^1,2,10,11^. Consistent with dopamine not being critical for aversion, Parkinson’s disease patients are known to exhibit paradoxical kinesia, a sudden ability to make quick and fluent movement in response to aversive stimuli^12-14^. Thus, we hypothesize that the other key neuromodulator ACh may play a critical role in signaling aversion that drives avoidance.

The ACh-releasing ChIs powerfully regulate striatal output via muscarinic receptors on striatal projection neurons, and indirectly through nicotinic receptors on GABAergic interneurons and on presynaptic terminals of cortical and subcortical inputs^15,16^. Studies using upregulation and downregulation of ChI activities have revealed their roles in sensorimotor integration, cognitive flexibility, and various aspects of reward learning^17-21^. ChIs are tonically active and fire at ∼2–10 Hz^22-24^. Their spiking is suppressed by sensory cues predicting both reward^25,26^ and aversive outcomes^27^, which are sometimes preceded by a brief excitation and often followed by a rebound^24,28,29^. As ChI suppression coincides with dopamine signaling, it has been hypothesized that ChI suppression defines a temporal window that permits learning^3,30-32^. However, recent studies show that acetylcholine suppression and dopamine signaling are temporally mismatched^33^, and ChI excitation is critical for obtaining reward requiring high effort^21^, supporting a more active role of ChIs in sensory processing and learning.

To test the role of ChIs in reinforcement learning, we performed nearly kilohertz cellular voltage imaging using high-performance genetically encoded voltage sensors SomArchon^34^ and ElectraOFF^35,36^ in mice during Pavlovian aversive and reward conditioning tasks. By recording sensory-evoked changes in subthreshold membrane voltage dynamics and suprathreshold spiking from individual ChIs, we probed ChI responses evoked by different behavioral events and the effect of learning. To further test the causal role of ChIs in behavior, we performed temporally precise optogenetic silencing of ChIs.

## Result

### Voltage imaging during aversive eye-blink conditioning and appetitive cued-reward conditioning tasks

To understand the role of ChIs in reinforcement learning, we performed cellular membrane voltage (Vm) imaging in head fixed mice during Pavlovian aversive and reward tasks (**Fig. 1A-E**). Briefly, we surgically implanted a custom imaging window coupled with an infusion cannula over the dorsal striatum and affixed a headplate for head fixation in ChAT-Cre mice (**Methods**). AAV9-FLEX-SomArchon-BFP, AAV9-FLEX-SomArchon-GFP or AAV9-DIO-ElectraOFF were infused into the dorsal striatum through the infusion cannula, transducing ChIs with the genetically encoded voltage indicator SomArchon or ElectraOFF. 2–4 weeks later, mice were habituated to head fixation and then recorded during the two conditioning tasks.

**Fig. 1.**
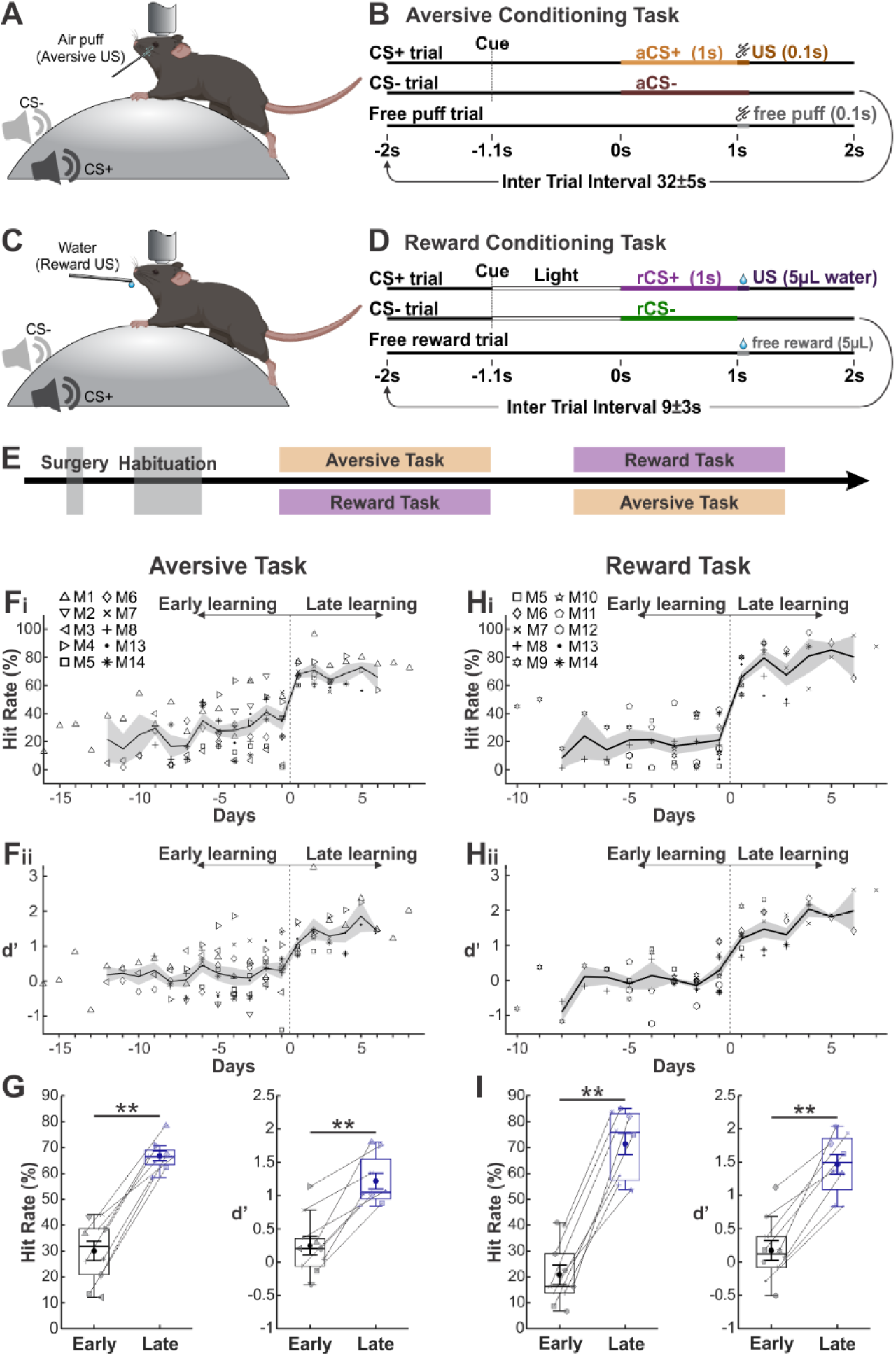
Experimental design and behavioral results. (**A**) Schematic of the experimental setup for the aversive task. The mouse icon was adapted from BioRender.com. (**B**) Illustration of the aversive task structure. Each trial is 4 s long, starting with a single sound click serving as the trial-start-cue. CS+ trials contain a sound (conditioned stimulus, aCS+, 1.1 s, orange), starting 1.1 s after trial-start-cue, which is followed by a 100 ms long air puff (unconditioned stimulus, US). CS-trials have a different sound (aCS-, 1.1 s, maroon) followed by nothing. Free puff trials only have puffs. Intertrial intervals are 32 ± 5 s. (**C**) Schematic of the setup for the reward task. (**D**) Illustration of the reward task structure. Like the aversive task, each trial is 4 s, starting with the same sound click as the trial-start-cue, which is followed by 1.1 s environmental white LED light until CS onset. CS+ trials have a sound (rCS+, 1.0 s, purple) followed by 5 µL water (US) accompanied by a 100 ms long auditory cue. CS- trials have a different sound (rCS-, 1.0 s, green) followed by nothing. Intertrial intervals are 9 ± 3 s. (**E**) Experimental timeline. (**F**) Hit rate and discriminability index (d′) across days in the aversive task, aligned to the first day when mice reached the behavioral learning criterion (day 0). Each dot represents a training session, and symbol shape indicates individual animals (n = 10 mice). Solid lines indicate the mean and shaded areas represent SEM. (**G**) Hit rate and d’ across animals during early (grey) versus late (blue) learning of the aversive task. Each symbol indicates the mean rate for an animal across all sessions within a learning stage, with lines connecting the same animals across learning stages (n = 8 mice). Boxes indicate the interquartile range (IQR; 25th–75th percentiles). Center lines are the median, and whiskers are the 1.5 × IQR. Mean ± SEM are overlaid on the box plots. Horizontal lines with asterisks indicate comparisons between learning stages using the Wilcoxon signed-rank test (Hit rate: p = 0.008; d’: p = 0.008). (**H**) Similar to (F) but for the reward task (n = 10 mice). (**I**) Similar to (G) but for the reward task (Wilcoxon signed-rank test, n = 8 mice. Hit rate: p = 0.008; d’: p = 0.008). * p < 0.05, ** p < 0.01, *** p < 0.001.

The aversive task was a classic eye-blink conditioning task, where mice were presented with one of two auditory conditioned stimuli (CS) for 1.1 s (*n* = 10 mice; **Fig. 1A-B, Methods**). 1 s after the onset of the positive CS (aversive CS+, aCS+), a 100 ms air puff unconditioned stimulus (US) was delivered to one eyelid. Following the negative CS (aversive CS-, aCS-), no air puff was delivered. Mouse behavior was video-recorded and analyzed using DeepLabCut ^37^ (**Fig. S1**). An eyeblink after aCS onset and before US onset was classified as conditioned response (CR+), and lack of blink during the same period was classified as no conditioned response (CR-). The reward task had a similar temporal structure, but with reward CS+ (rCS+) followed by a water reward (US) and reward CS- (rCS-) followed by nothing (*n* = 10 mice; **Fig. 1C-D**). Two or more consecutive licks within 1 s after rCS onset were classified as CR+, and zero licks during the same time window were classified as CR-. Six animals were trained on both tasks, and the order of the aversive and reward task was randomized across these animals.

Behavioral performance was quantified by hit rate (CS+ CR+ trials divided by all CS+ trials) and d’ (z-scored hit rate minus z-scored false alarm rate, calculated as the ratio of CS- CR+ trials over all CS- trials). 8 mice learned the aversive task after 638 ± 74 trials (mean ± SEM, *n* = 8 mice; **Fig. 1F**), defined as having reached a behavioral threshold of hit rate > 50% and d’ > 0.7. During late learning, defined as sessions after reaching the behavioral threshold, the hit rate and d’ were significantly higher than during early learning before reaching the threshold (**Fig. 1G**). Similarly, 8 mice learned the reward task over 358 ± 82 trials (*n* = 8 mice; **Fig. 1H**), with significantly increased hit rate and d’ in late learning than in early learning (**Fig. 1I**). The 4 mice that did not reach behavioral threshold were only included in the analysis of the early learning stage.

### ChIs exhibit prominent immediate excitation followed by delayed inhibition to behaviorally relevant events during both tasks

To capture ChI dynamics throughout learning, voltage imaging was performed on the first day of training and during most training days. Animals expressing SomArchon were trained for 60 trials per day to minimize photobleaching, and animals expressing the more photostable ElectraOFF were trained for 100 trials per day. Recorded image stacks were motion corrected, segmented to extract fluorescence at the soma, and then processed to obtain Vm and spikes (**Methods**). We did not notice any difference across SomArchon and ElectraOFF recordings, and thus all following analyses were performed on combined datasets.

We noted that many ChIs exhibited a robust and transient Vm depolarization and firing rate (FR) increase, following the trial-start-cue and both CSs in both tasks (**Fig. 2, Fig. S2, Fig. S3**). The transient excitation generally had a short latency of ∼14–23 ms (**Table S2**) and peaked around 35–100 ms (defined as immediate window), which was often followed by a prolonged Vm hyperpolarization and spike suppression that peaked ∼250–450 ms. While the aversive US evoked similar immediate excitation as the cue and CSs, the reward US evoked weaker and delayed activation occurring around 100–600 ms after US onset (**Fig. 2B, D**).

**Fig. 2.**
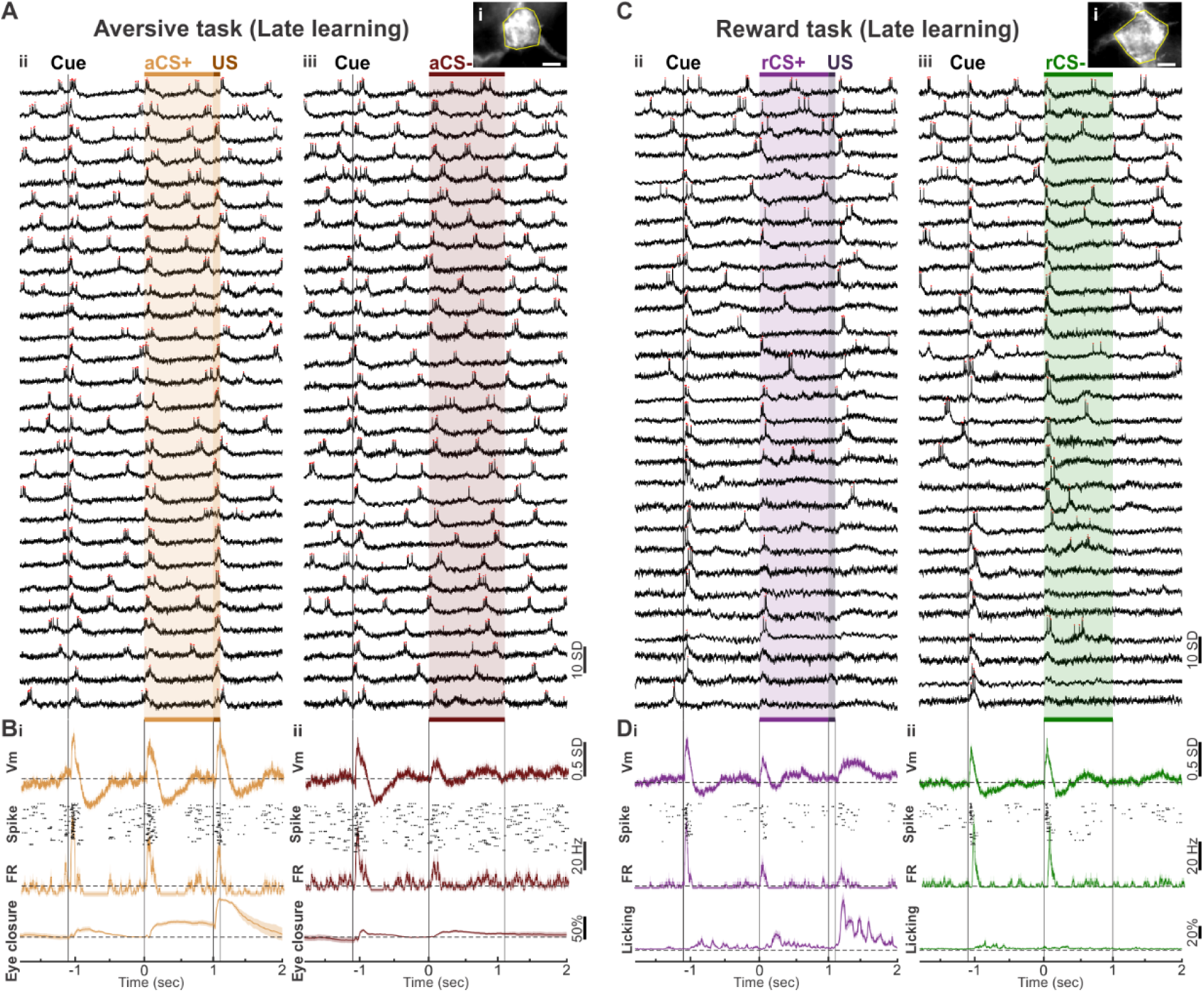
ChIs exhibit prominent immediate excitation followed by delayed inhibition to behaviorally relevant events during both tasks. (**A, B**) An example recording session from one ElectraOFF expressing ChI during the aversive task in late learning. (**Ai**) Upper right: ElectraOFF fluorescence image, averaged across all image frames during one trial. Yellow outline indicates the ROI selected as the ChI soma; scale bar, 10 μm. (**Aii-iii**) Vm traces (black) across example trials aligned to CS onset. Red dots indicate spikes. (**Aii**) Aversive CS+ trials. aCS+ duration is highlighted by orange shading. US: dark brown shading. (**Aiii**) Aversive CS- trials. aCS- duration is highlighted by maroon shading. Trial-start-cue is labelled as vertical dashed line. (**B**) Vm, FR (shown with 6 ms smoothing window) overlaid with spike raster, and eyelid movement across all aCS+ (**Bi**, orange: n = 28 trials) and aCS- (**Bii,** maroon: n = 28 trials) trials. Vm, FR, and eyelid movement were baseline-subtracted using the corresponding mean during -100 to 0 ms pre-CS baseline for each trial. Dashed lines are at 0. Solid lines indicate the mean and shaded areas represent SEM. (**C**) Similar to (A), but for an example session during the reward task. (**Ci**) An ElectraOFF expressing ChI. (**Cii**) Reward CS+ (purple shading) is followed by water reward (dark purple); (**Ciii**) Reward CS- (green shading) is followed by nothing. (**D**) Similar to (B), but with the behavior shown as lick movement. (**Di**) Across rCS+ trials (purple: n = 29 trials); (**Dii**) Across rCS-trials (green: n = 31 trials).

### Aversive learning augments ChI immediate excitation and delayed inhibition to aCS+ predicting punishment, but not to aCS- predicting no punishment

To characterize the immediate response of individual ChIs during aversive learning, we first determined whether a neuron was modulated by each task-related event. Specifically, the mean Vm or FR during the immediate window was compared to the pre-event baseline (-100 to 0 ms) across trials for each neuron. We found that >94% of ChIs were activated (Vm depolarization and FR increase) by the trial-start-cue in both learning stages, and 60–78% were also activated by the US (**Table S2**). However, aCS+ and aCS- evoked responses varied across neurons in early learning **(Fig. 3A, C)**, activating 47–66% of neurons (**Table S2**). In late learning, many ChIs exhibited more pronounced immediate excitation to aCS+ (**Fig. 3Bi, Di**), with 92.6% being Vm activated and 78% FR activated, significantly higher than during early learning (*p* = 0.016 and 0.020 respectively, Fisher’s exact test; **Table S2**). In contrast, aCS- evoked Vm and FR responses during late learning was similar to early learning (**Fig. 3Bii, Dii; Table S2**). Thus, aversive learning selectively enhances the fraction of ChIs exhibiting immediate Vm depolarization and FR increase to aCS+, without affecting their response to aCS-.

**Fig. 3.**
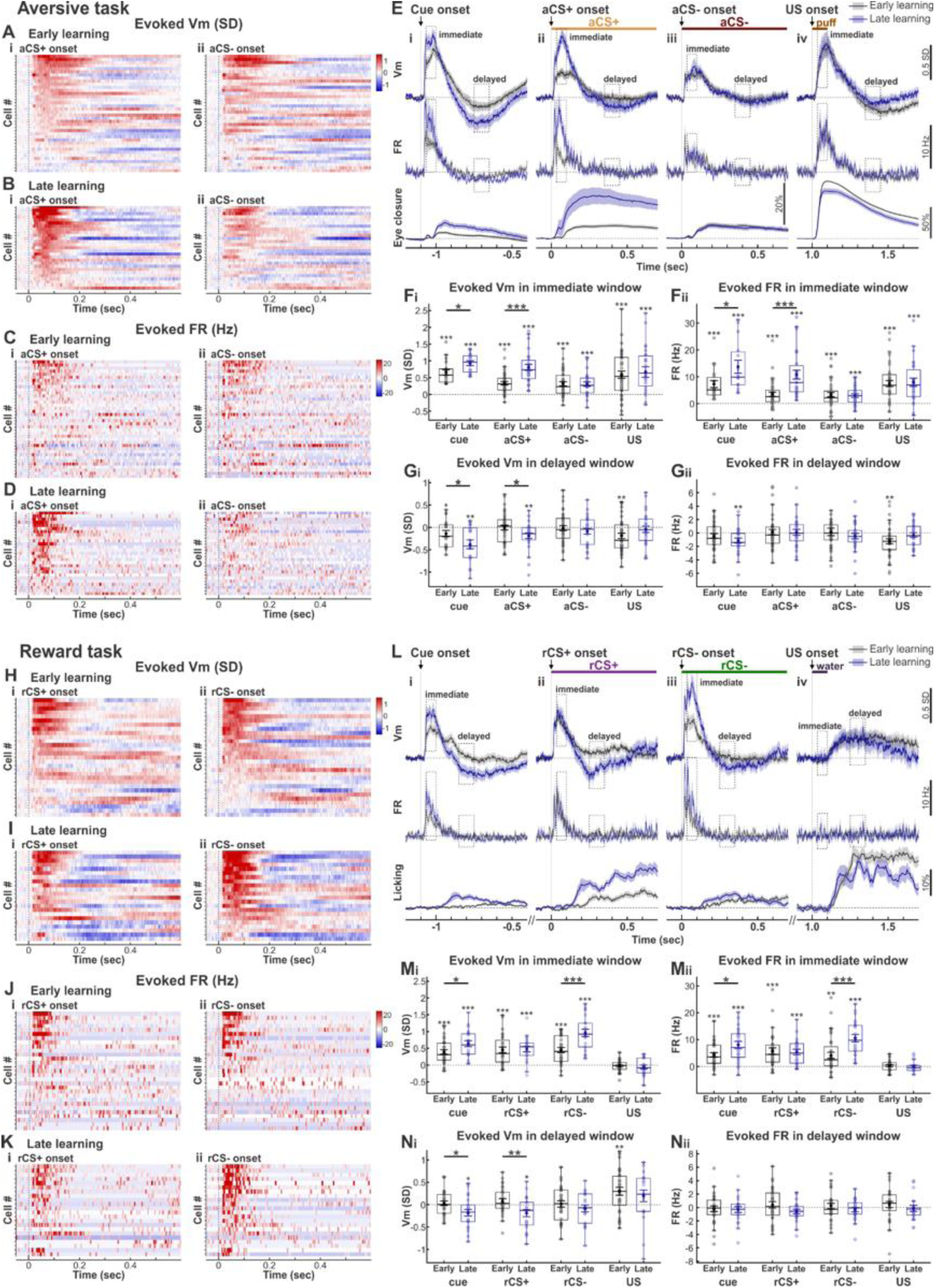
ChI responses to CS predicting negative outcomes are selectively augmented by learning in both tasks. (**A-G**) ChI responses during the aversive task. **(A)** Heatmaps of the evoked Vm by aversive CS+ (Ai) and aversive CS- (Aii) across all neurons in early learning. Each row represents one neuron (n = 38 neurons from 10 mice). Evoked Vm was computed as the trial-wise mean of the baseline subtracted Vm, with the baseline defined as the mean during the -100 to 0 ms pre-event period for each trial. Neurons are sorted by the amplitude of the evoked Vm by aCS+ during the immediate window. (**B**) Similar to (A) but for late learning (n = 27 neurons from 8 mice). (**C-D**) Similar to (A-B) but for FR. (**E**) Population Vm, FR and eye closure across recorded neurons aligned to trial-start-cue (Ei), aCS+ (Eii), aCS- (Eiii), and US (Eiv) during early (grey) and late (blue) learning. Dashed lines are at 0. Solid lines indicate mean and shaded areas represent SEM (early learning, aligned to cue n = 19 neurons from 5 mice, aligned to aCS+/aCS- and US n = 38 neurons from 10 mice; late learning, aligned to cue n = 17 neurons from 4 mice, aligned to aCS+/aCS- and US n = 27 neurons from 8 mice). Immediate and delayed windows are highlighted by the dashed boxes. (**F-G**) Event-evoked Vm **(Fi, Gi)** and FR (**Fii, Gii**) during the immediate (**F**) and delayed (**G**) windows for trial-start-cue, aCS+, aCS- and US. Boxes indicate the interquartile range (IQR; 25th–75th percentiles) across neurons. Center lines are the median, and whiskers are the 1.5 × IQR. Mean ± SEM are overlaid on the box plots. Asterisks above boxes indicate comparisons to baseline using the Wilcoxon signed-rank test; horizontal lines with asterisks indicate comparisons between learning stages using the Mann-Whitney U test (see **Table S5** for statistical details). (**H-N**) Similar to (A-G) but for the reward task (early learning, n = 29 neurons from 10 mice; late learning, n = 22 neurons from 8 mice, US, n = 19 due to motion exclusion). In (H-K), neurons are sorted by the amplitude of the evoked Vm by rCS- during the immediate window. In (L), behavior is shown as licking movement instead of eye closure. * p < 0.05, ** p < 0.01, *** p < 0.001.

We then evaluated population evoked responses by first normalizing each neuron’s Vm and FR by subtracting the pre-event baseline (**Fig. 3E**). Like most individual neurons, ChI population exhibited robust and significant Vm depolarization and FR increase during the immediate window following all task-relevant events (trial-start-cue, aCSs, and US) (**Fig. 3F**). Furthermore, learning selectively increased the immediate excitation to aCS+, without altering the response to aCS-(**Fig. 3Fi-Fii**). Additionally, consistent with the characteristic shift of response to earlier cues in Pavlovian conditioning, the immediate excitation evoked by trial-start-cue was enhanced by learning, whereas US-evoked excitation remained unchanged throughout learning (**Fig. 3F**). Finally, we examined ChI responses to free air puffs in a subset of neurons in both early and late learning, and confirmed that free puff evoked greater or comparable excitation to all other behaviorally relevant events (**Fig. S4B-E**).

Across ChI population, the immediate excitation was followed by a large Vm hyperpolarization peaking around 350–450 ms (delayed window). We found that learning significantly augmented the evoked hyperpolarization following the trial-start-cue and aCS+, but not aCS- (**Fig. 3Gi**). However, the strong Vm hyperpolarization was not accompanied by significant FR suppression (**Fig. 3Gii**), likely due to the generally low ChI basal FR (2.89 ± 0.14 Hz, *n* = 28 neurons).

As ChIs may be modulated by movement, we aligned ChI Vm and spiking to the onset of eye closure and detected no obvious temporal relationship between eye closure and ChI responses (**Fig. S5**). Further analysis of aCS-evoked excitation between CR+ and CR- trials revealed no difference in either aCS+ or aCS- trials during late learning (**Fig. S5A-C, F-H**), again confirming that ChI responses to aCSs were not due to movement. Together, these results demonstrate that ChIs are robustly activated by behaviorally salient stimuli, often followed by prolonged Vm hyperpolarization. ChI responses are shaped by aversive learning, exhibiting selective augmentation of both immediate excitatory and delayed inhibitory responses to aCS+ predicting aversive eye-puff, but not to aCS- predicting no punishment.

### Reward learning selectively augments ChI immediate excitation to rCS- predicting reward omission, but not to rCS+ predicting reward, and has the opposite effect on delayed inhibition

During reward learning, most neurons (>72%) were activated (Vm depolarization and FR increase) by trail-start-cue during both stages and few (<10%) showed immediate activation to the US (**Table S2**). As in the aversive task, most neurons showed similar Vm and FR changes to the two rCSs in early learning (**Fig. 3H, J**). However, in late learning, ChI exhibited stronger excitation to rCS- but not to rCS+ (**Fig. 3I, K**), with a greater fraction of ChIs being excited by rCS-during late learning than early learning (*p* = 0.039 and 0.0098, Vm and FR respectively; **Table S2**). In contrast, the fraction of Vm and FR modulated neurons by rCS+ remained unchanged (**Table S2**). Thus, reward learning selectively enhances the fraction of ChIs activated by rCS-, without affecting the response to rCS+.

Further analysis of the ChI population responses revealed robust Vm depolarization and FR increase following the trial-start-cue and both rCSs, peaking around 35–100 ms (immediate window), but weak and delayed activation by the US, in both early and late learning (**Fig. 3L, M**). Furthermore, the immediate excitation to rCS- were significantly higher in late learning than early learning for both Vm depolarization and FR increase, whereas responses to rCS+ remained unchanged throughout learning (**Fig. 3Mi-Mii**). Interestingly, ChIs were robustly activated by free reward, in sharp contrast to the weak responses following the US (**Fig. S6A**). Immediate excitation to free reward was similar to the trial-start-cue and rCS+, and significantly larger than that evoked by the US (**Fig. S6B, C**), suggesting that ChIs respond more strongly to salient events than to predicted reward.

The delayed inhibition following trial-start-cue and rCS+ was minimal in early learning, which emerged in late learning with significant Vm hyperpolarization around 250–350 ms (delayed window) (**Fig. 3Mi, Ni**). In contrast, delayed inhibition remained minimal following rCS- throughout learning, even though the evoked excitation augmented by learning (**Fig. 3Mi, Ni**). As ChIs exhibited a low basal FR (2.79 ± 0.16 Hz, *n* = 45 neurons), and FR suppression was minimal during the delayed window (**Fig. 3Nii**). Thus, reward learning amplifies ChI immediate Vm depolarization and FR increase only to rCS-, and enhances delayed Vm hyperpolarization to trial-start-cue and rCS+.

Additionally, aligning Vm and spiking to lick onset revealed no obvious temporal relationship between licking and evoked responses (**Fig. S7**). Furthermore, ChI responses were similar between CR+ and CR- trials following either rCS+ or rCS- (**Fig. S7A-C, F-H**), and exhibited no Vm or FR changes around lick onset (**Fig. S7D-E, I-J**). Overall, these results demonstrate that ChIs are reliably activated by behaviorally salient stimuli during reward learning. Learning selectively augments immediate ChI excitation to the rCS-, which predicting unfavorable reward omission, without affecting the response to the rCS+ predicting reward. While immediate excitation evoked by rCS+ was not altered by reward learning, delayed inhibition was augmented, suggesting that delayed inhibition is likely driven by network inputs that are shaped by reward learning.

### ChIs exhibit greater immediate excitation, but not delayed inhibition, to CS predicting negative outcomes in both tasks during late learning

After observing that learning selectively augments immediate excitation to CS predicting negative outcomes in both tasks (**Fig. 3**), we directly compared the response evoked by CS+ versus CS-in each neuron. In early learning, few ChIs showed a significant difference in their Vm (∼12%) and FR (∼6%) response to the two CSs in both tasks (**Table S3**). In late learning, a similar fraction (∼52%) of neurons exhibited greater Vm responses to CSs predicting negative outcome than those predicting positive outcome (**Table S3**). However, a greater fraction of neurons showed augmented FR response to aCS+ than rCS- (*p* = 0.019; **Table S3**), suggesting the population ChI spiking output contributes more significantly to punishment than reward omission, even though the input strengths are similar.

To further assess the temporal dynamics of the differential response to the two CSs, we computed the difference in the evoked Vm (ΔVm, CS+ minus CS-) and FR (ΔFR) in each ChI (**Fig. 4Ai,ii-Di,ii**). Across the ChI population, ΔVm and ΔFR remained flat for the entire period from CS onset to US onset in both tasks during early learning, again confirming a lack of discrimination between CS+ and CS- at the population level (**Fig. 4Aiii-Diii, E-H**). However, during late learning, aCS+ evoked population response was greater than that of aCS- in the aversive task (**Fig. S8A, B, E, F**), leading to significantly positive ΔVm and ΔFR during 15–190 ms and 10–115 ms post aCS onset respectively (**Fig. 4Aiii, Biii, Ei, Fi**). In the reward task, rCS- evoked greater immediate excitation than rCS+ (**Fig. S8C, D, G, H**), with ΔVm and ΔFR being significantly negative during 12–170 ms and 14–100 ms post rCS onset respectively (**Fig. 4Ciii, Diii, Eii, Fii**). Consistent with some neurons also exhibited stronger delayed inhibition to rCS+ than to rCS- in the reward task, we observed an additional negative deflection during 220–290 ms post rCS onset (**Fig. 4Cii, Ciii**).

**Fig. 4.**
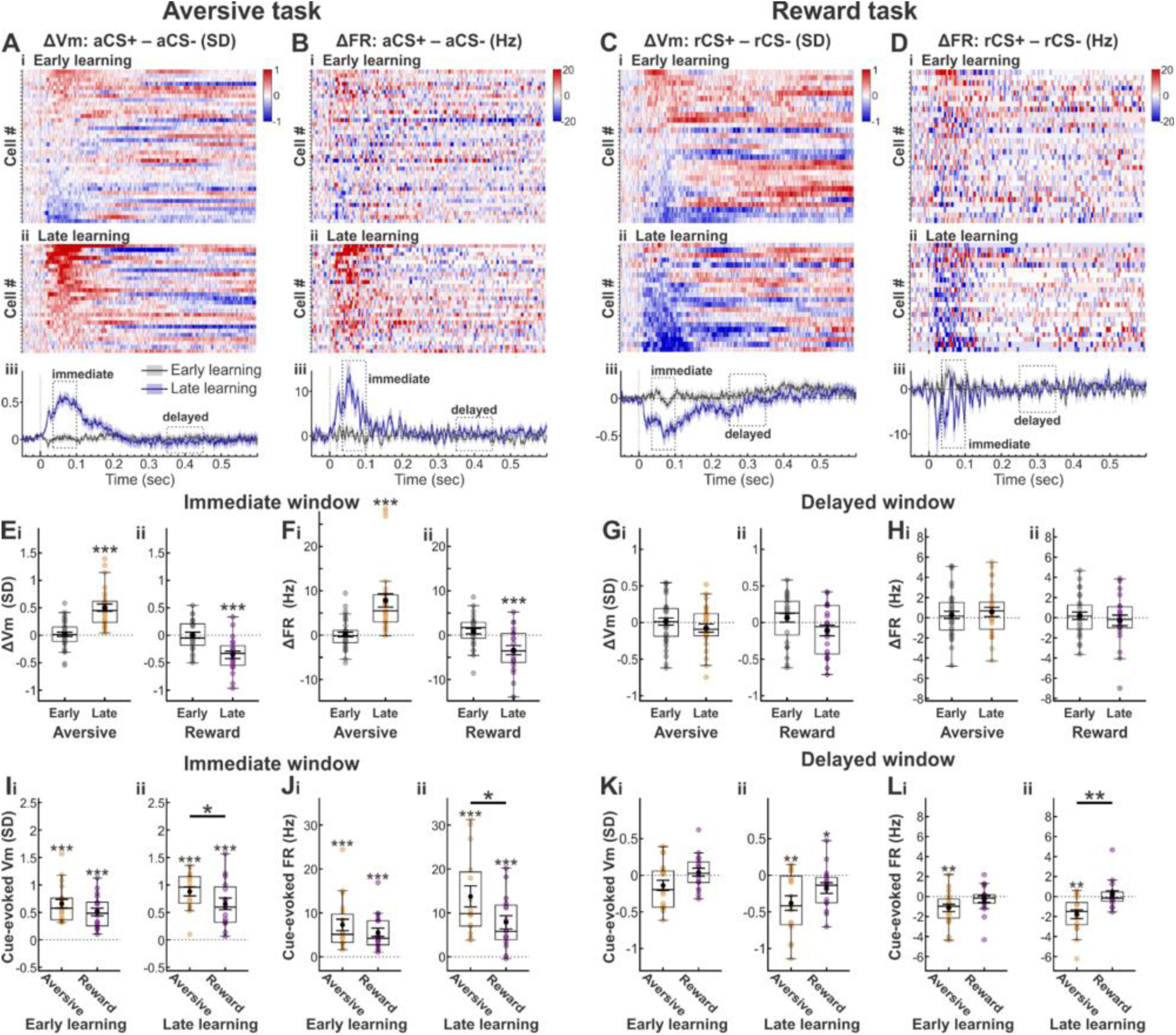
ChI Vm and spiking exhibit greater response to CSs predicting negative outcomes. (**A**) Heatmaps of the differential Vm (ΔVm) evoked by aCS+ minus aCS- trials across neurons during early (Ai, n = 38 neurons from 10 mice) and late (Aii, n = 27 neurons from 8 mice) learning. Each row represents a neuron. Neurons are sorted by the amplitude of ΔVm during the immediate window. (**Aiii**) Population ΔVm across all neurons during early (grey) and late (blue) learning. Solid lines indicate mean and shaded areas represent SEM. Immediate and delayed windows are highlighted by dashed boxes. (**B**) Similar to (A) but for differential FR (ΔFR). (**C-D**) similar to (A-B) but for the reward task (n = 29 neurons from 10 mice for early learning, n = 22 neurons from 8 mice for late learning). (**E-H**) ΔVm (E, G) and ΔFR (F, H) during the immediate (E, F) and delayed (G, H) window in the two tasks. Boxes indicate the interquartile range (IQR; 25th–75th percentiles) across neurons. Center lines are the median, and whiskers are the 1.5 × IQR. Mean ± SEM are overlaid on the box plots. Asterisks above boxes indicate comparisons to zero using the Wilcoxon signed-rank test (see **Table S8** for statistical details). (**I-L**) Similar to E-H, but for evoked response by trial-start-cue (Early learning, n = 19 neurons from 4 mice for aversive; n = 18 neurons from 7 mice for reward. Late learning, n = 17 neurons from 4 mice for aversive; n = 16 neurons from 5 mice for reward). Asterisks above boxes indicate comparisons to baseline using the Wilcoxon signed-rank test; horizontal lines with asterisks indicate comparisons between two tasks using the Mann-Whitney U test (see **Table S10** for statistical details). * p < 0.05, ** p < 0.01, *** p < 0.001.

While CS+ evoked delayed hyperpolarization was slightly augmented by learning in both aversive and reward tasks (**Fig. 3Gi, Ni**), the fraction of neurons showing differential response to the two CSs were identical throughout learning (**Table S3**), consistent with the population ΔVm and ΔFR being around baseline levels during the delayed window (**Fig. 4Aiii-Diii, G, H, Fig. S8A-H**). Together, these results indicate that immediate ChI excitation preferentially signals negative outcome, either punishment or reward omission. In contrast, delayed inhibition does not contribute to outcome discrimination.

To further probe ChI response preference between aversive versus reward conditions, we compared the magnitude of responses evoked by identical trial-start-cue between the two tasks. In early learning, the trial-start-cue evoked immediate and delayed responses were comparable between the two tasks (**Fig. 4Ii-Li, Fig. S8I, K**). However, in late learning, trial-start-cue evoked immediate Vm depolarization and FR increase were greater in the aversive task than in the reward task (**Fig. 4Iii, Jii, Fig. S8J, L**), suggesting context-dependent modulation of excitatory inputs to ChIs.

Furthermore, in late learning, cue-evoked Vm hyperpolarization showed no significant difference (**Fig. 4Kii, Fig. S8J**), unlike immediate Vm depolarization, though late FR suppression was stronger in the aversive task than in the reward task (**Fig. 4Lii, Fig. S7L**), indicating different network contributions to the excitatory versus inhibitory ChI responses. To further probe this possibility, we examined the correlation between immediate and delayed responses, and detected little correlation (**Fig. S8M-N**). Together, these results highlight that ChI dynamics are differentially modulated during the immediate and delayed phases, suggesting different inputs drive these temporally distinct responses in both tasks.

### Comparing the same neurons recorded in both tasks further confirms that ChIs signal negative outcomes, and reveals task-preferential activation of inputs to individual ChIs

With the ability to visualize cell morphology and surrounding tissue and vasculature, it is possible to record from the same neurons across days (**Fig. 5A**). We recorded from 10 ChIs in both tasks during the late learning (**Fig. 5B-I**), with counterbalanced behavioral task orders across mice (**Table S4**). The CS and trial-start-cue evoked responses in these 10 neurons during both tasks are consistent with those observed across the larger population, further confirming that ChIs signal negative outcomes (**Fig. 5J-M**), and exhibit enhanced excitatory response in the aversive context (**Fig. 5O-R**). Next, we compared the same neuron’s ability in discriminating CSs across tasks, and found that more ChIs showed CS discrimination in the aversive task (9 of 10 neurons) than in the reward task (4 neurons), with 2 showing discrimination in both tasks. Evoked FR increase showed a similar trend in discriminating the two CSs, though with fewer neurons passing the statistical threshold due to sparse firing (**Table S4**).

**Fig. 5.**
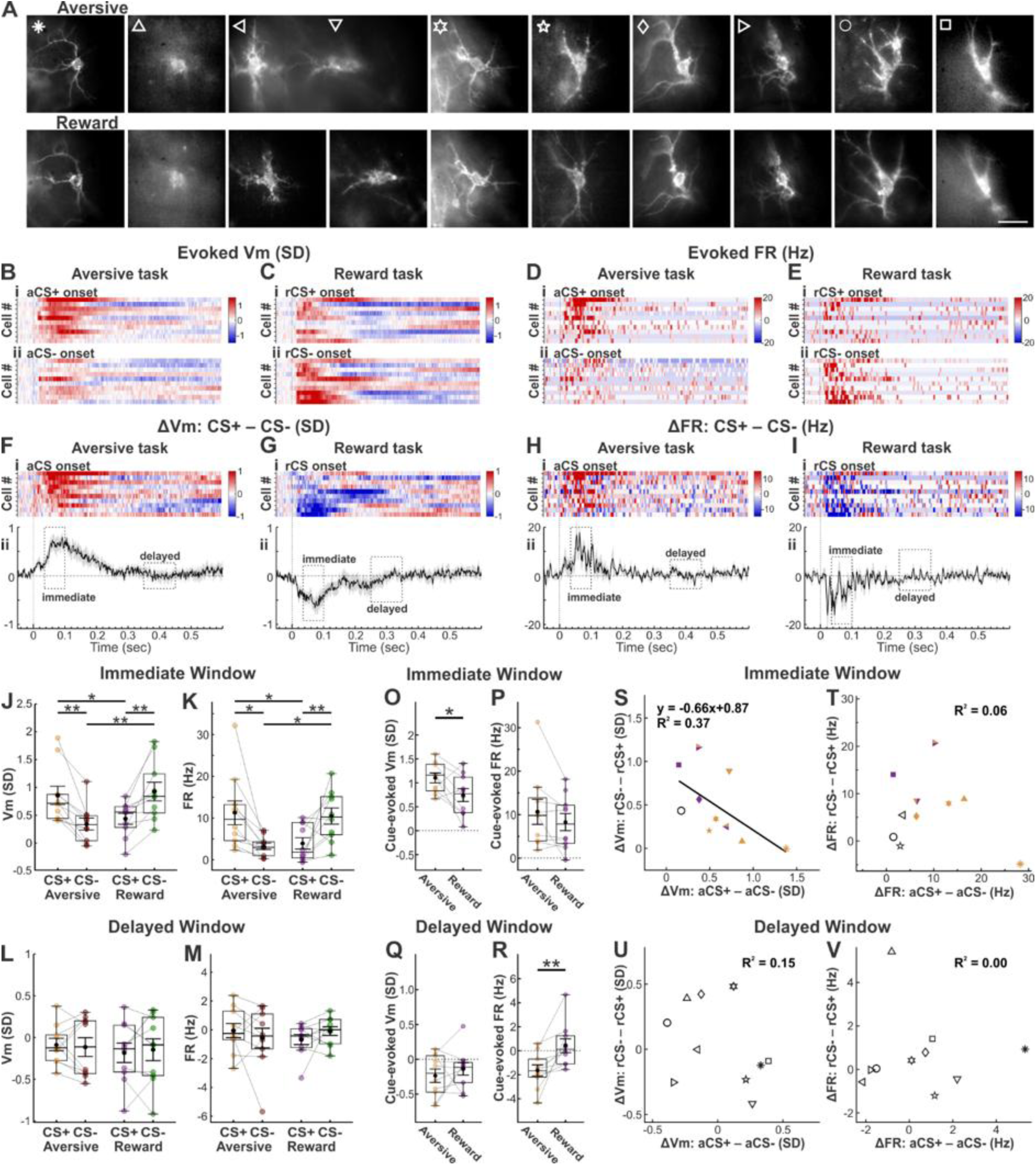
The same neurons recorded in both tasks reveal task-preferential engagement of distinct ChIs. (**A**) A single frame fluorescence images of the same 10 ChIs from 5 mice recorded during both the aversive (top) and reward (bottom) tasks in late learning (scale bar, 10 μm). Neurons are sorted from left to right by the amplitude of the evoked Vm difference during the immediate window (as sorted in Fi). (**B**) Heatmaps of evoked Vm by aCS+ (Bi) and aCS- (Bii) in the aversive task. Each row represents one neuron. Neurons are sorted in the same order as in Fi. (**C**) Similar to (B) but for the reward task, with the same neuron sorting. (**D-E**) Similar to (B-C) but for FR. (**F**) Heatmaps of the differential Vm response (ΔVm) evoked by CS+ minus CS- trials (Fi), and population ΔVm (Fii) across 10 neurons in the aversive task. Solid lines indicate mean and shaded areas represent SEM. Immediate and delayed windows are highlighted by dashed boxes. (**G**) Similar to (F) but for the reward task, with neurons in Gi sorted in the same order as in Fi (**H-I**) Similar to (F-G) but for ΔFR. (**J-K**) Comparisons of the evoked Vm (J) and FR (K) during the immediate window by aCS+ (orange), aCS- (maroon), rCS+ (purple) and rCS- (green). Boxes indicate the interquartile range (IQR; 25th–75th percentiles) across neurons. Center lines are the median, and whiskers are the 1.5 × IQR. Mean ± SEM are overlaid on the box plots. Horizontal lines with asterisks indicate comparisons between conditions using the Wilcoxon signed-rank test (see **Table S11** for statistical details). (**L-M**) Similar to (J-K) but for the delayed window. (**O-R**) Similar to (J-M) but for evoked responses by trial-start-cue across aversive and reward tasks (n = 9). Horizontal lines with asterisks indicate pairwise comparisons between tasks using the Wilcoxon signed-rank test (see **Table S11** for statistical details). (**S**) Scatter plot of ΔVm in the aversive (aCS+ minus aCS-) versus reward (rCS- minus rCS+) tasks across neurons. Solid line indicates linear regression fits, with equation and R^2^ shown in panel. Symbol shapes correspond to the neuron shown in (A). Orange-filled symbols indicate neurons that discriminated the CSs only in the aversive task, purple-filled symbols indicate neurons that discriminated only in the reward task, and dual-colored symbols indicate neurons that discriminated in both tasks (Mann-Whitney U tests, see **Table S4** for statistical details). (**T**) Similar to (S) but for ΔFR. (**U-V**) Similar to (S-T) but for delayed window. Only R² values are shown due to weak linear fits. * p < 0.05, ** p < 0.01, *** p < 0.001.

To explore the CS discrimination ability of the same neurons between aversive versus reward contexts, we examined the difference in the evoked response by negative versus positive predicting CSs (relative ΔVm and ΔFR). We detected a significant and reasonably strong inverse correlation of the immediate ΔVm between tasks, suggesting that neurons showing stronger response to negative outcome predicting CS in one task tend to show weaker response in the other (**Fig. 5S**). In contrast, delayed ΔVm showed no correlative relationship (**Fig. 5U**), consistent with the delayed inhibition being insensitive to task-specific outcome predictions. The relative ΔFR in the two tasks showed no relationship (**Fig. 5T, V**), again likely due to sparse firing. Together, these results support diverse and task-dependent activation of inputs to individual ChIs.

### Optogenetic silencing of ChI immediate excitation interferes with the avoidance behavior, but not the reward-seeking behavior

After observing a selective augmentation of immediate excitatory response to CSs predicting negative outcomes, we next tested the causal role of this immediate excitation in behavioral performance. AAV9-FLEX-ArchT-GFP was infused into the dorsal striatum of ChAT-Cre mice to selectively express the optogenetic silencer ArchT in ChIs (**Methods**). Six mice were trained on the two tasks for 100 trials per day. Five reached the late learning stage in the aversive task after 920 ± 116 training trials (mean ± SEM, *n* = 5 mice; hit rate = 59.0 ± 3.2% and d’ = 0.86 ± 0.06), and five reached the late learning stage in the reward task after 440 ± 51 training trials (*n* = 5 mice; hit rate = 87.2 ± 3.3% and d’ = 1.79 ± 0.15). Once reaching late learning stage, we tested the effect of ChI silencing either during the immediate (0–300 ms after CS onset) or the delayed (300–600 ms) windows (**Fig. 3**). Each recording session contained 4 types of trials, with (ArchT-ON) or without silencing (ArchT-OFF), following CS+ or CS- (**Fig. 6A, E, I, M**). The four trial types were pseudorandomized, with immediate and delayed silencing performed on different days.

**Fig. 6.**
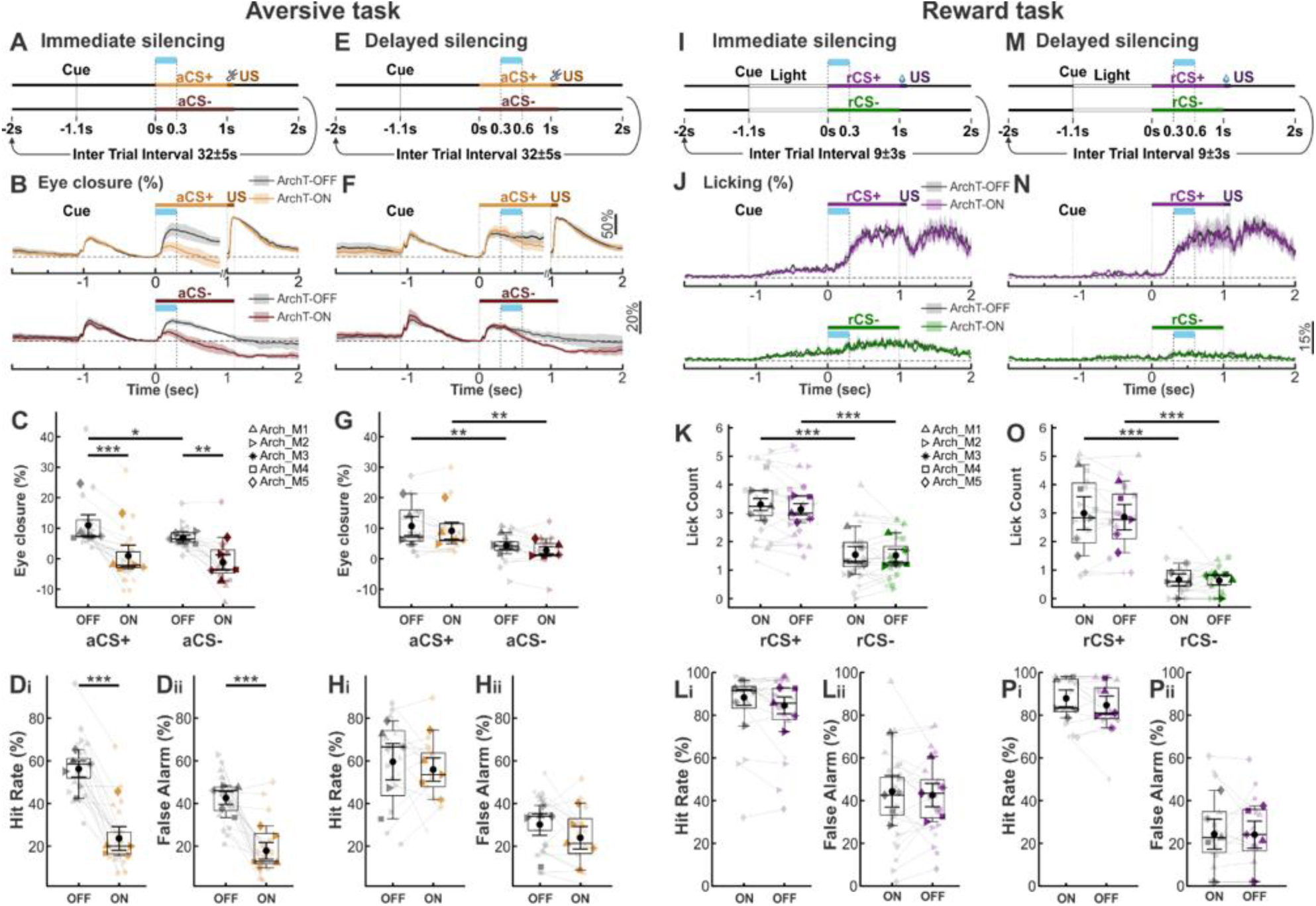
Optogenetic silencing of ChIs impairs performance in the aversive task but not the reward task post learning. (**A-H**) Optogenetic silencing of ChIs during late learning in the aversive task. (**A**) Experimental protocol. Immediate silencing occurred during 0–300 ms after CS onset. (**B**) Eye closure across ArchT-OFF (grey) and ArchT-ON (colored, orange for aCS+, maroon for aCS-) trials. Eye closure was normalized by subtracting the mean during the -100 to 0 ms before CS onset. Dashed lines are at 0. Solid lines indicate mean and shaded areas represent SEM (n = 5 mice). (**C**) Box plots of eye closure from aCS onset to US onset during ArchT-OFF versus ArchT-ON trials. Boxes indicate the interquartile range (IQR; 25th–75th percentiles) across neurons. Center lines are the median, and whiskers are the 1.5 × IQR. Mean ± SEM are overlaid on the box plots. Horizontal lines with asterisks indicate comparisons between conditions using the linear mixed-effects model (see **Table S12** for statistical details). Semi-transparent dots represent individual sessions with lines connecting the ArchT-OFF and ArchT-ON conditions within the same session, while solid dots indicate the session-wise mean for each mouse. Symbols indicate different mice. (**D**) Hit rate (Di) and false alarm (Dii) in ArchT-OFF (grey) and ArchT-ON (orange) trials. (**E-H**) Similar to (A-D) but for ChI delayed silencing that occurred during 300–600 ms after CS onset. **(I-P)** Optogenetic silencing of ChIs during late learning in the reward task (n = 5 mice). (**I-J**) Similar to (A-B) but for the reward task, with purple for rCS+, green for rCS-. (**K**) Box plots of lick numbers from CS onset to US onset with ChI immediate silencing (linear mixed-effects model; see **Table S12** for statistical details). (**L**) Box plots of hit rate (Li) and false alarm (Lii) in ArchT-OFF (grey) and ArchT-ON (purple) trials. (**M-P**) Similar to (I-L) but for ChI delayed silencing. * p < 0.05, ** p < 0.01, *** p < 0.001.

During the aversive task, immediate silencing significantly reduced eye closure following both aCS+ and aCS- and eliminated their difference (**Fig. 6B, C**), and decreased hit rate and false alarm rate (**Fig. 6B-D**). In contrast, delayed silencing produced no measurable behavioral effects (**Fig. 6E-H**), suggesting that immediate ChI excitation, but not delayed inhibition, is necessary for appropriate avoidance response. During the reward task, neither immediate nor delayed silencing produced any behavioral effects measured as lick count, hit rate and false alarm rate to either rCS+ or rCS-, and licking response to rCS+ remained higher than rCS- across all conditions as expected (**Fig. 6J-K, N-P**). The selective impact of ChI optogenetic silencing on behavioral performance in the aversive task, but not in the reward task, indicates that immediate ChI excitation is critical for avoidance behavior, but not reward-seeking behavior.

## Discussion

Using precision cellular voltage imaging in behaving mice, we examined subthreshold Vm and suprathreshold spiking of individual genetically defined cholinergic interneurons (ChIs) in the dorsal striatum during Pavlovian aversive and reward conditioning tasks. We found that ChIs exhibit robust Vm depolarization and increased spiking in response to task-relevant auditory stimuli with a short latency, followed by prolonged Vm hyperpolarization and spike suppression. Reinforcement learning selectively augments the immediate excitatory response evoked by conditioned stimulus (CS) predicting negative outcomes, either air puff or reward omission, without affecting that evoked by other CSs. While the delayed inhibition is also strengthened by learning, it does not distinguish between CSs. Finally, brief optogenetic silencing of ChIs following CS onset confirmed that immediate ChI excitation is required for avoidance behavior but not for reward-seeking. Together, these results demonstrate that precisely timed ChI activation signals aversion that is important for avoidance behavior.

### ChI immediate excitation is selectively augmented by learning and drives avoidance

Through near kilohertz voltage imaging, we reveal a rapid excitatory response following auditory stimuli with a remarkably short latency of ∼14–23 ms in most neurons. After this rapid onset, the excitation evoked by different CSs diverged in late learning, reaching maximum difference ∼35–100 ms after CS onset. This divergence is due to learning-dependent selective augmentation of the evoked excitation by CSs predicting negative outcomes. Furthermore, the identical trial-start-cue evoked greater augmentation of the immediate excitation in aversive learning than reward learning, suggesting a context-dependent increase in ChI excitatory responses to the same sensory events. As ChI excitation has been predominantly attributed to cortical^38,39^ and thalamic^40,41^ glutamatergic inputs, the difference in evoked Vm depolarization likely reflects learning-dependent augmentation of excitatory synaptic inputs onto ChIs, particularly those from the amygdala-prefrontal pathways known to be involved in aversive learning^42,43^. In addition to similar increase in synaptic response captured by Vm depolarization, learning led to a greater FR rate increase to aCS+ than rCS-. Thus, similar input strengthens captured by Vm depolarization led to a greater population ChI output to evidence predicting punishment than reward omission. Silencing of this immediate excitation using optogenetics selectively interfered with appropriate avoidance behavior in the aversive task, without affecting reward-seeking behavior, suggesting that ChI activity drives avoidance.

### Delayed inhibition is strengthened by learning

The immediate ChI excitation is followed by prolonged inhibition that peaked around 300 ms after stimulus onset. The delayed inhibition has been postulated to open a temporal window for DA-dependent plasticity to occur^44^. However, we observed that the two CSs evoked similar delayed inhibition regardless of task or learning stage, and silencing ChIs during the 300–600 ms after CS onset failed to alter behavioral performance in either task. Thus, while delayed inhibition in ChIs are shaped by learning, it does not drive avoidance or approach.

DA plays an important role in ChI delayed inhibition^45-47^. Aosaki et al. showed that ∼17% of extracellularly recorded ChIs (tonically active neurons) exhibited response suppression to CS before conditioning, which increased to 73% after reward learning, and DA depletion reduced the fraction back to ∼19%^26^. Consistent with these findings, we found that delayed inhibition following rCS+ was robustly augmented by reward learning. However, we also observed a significant strengthening following aCS+ by aversive learning, suggesting that various cellular and circuit mechanisms beyond DA are involved. At the cellular level, learning-mediated changes in ion channels underlying ChI intrinsic Vm delta rhythmicity (∼1–4 Hz) likely contribute to the augmented inhibition at ∼300 ms (3.3 Hz) during late learning. For example, learning-mediated increase in excitation can elevate Ca^2+^ influx during spiking, leading to greater hyperpolarization via K^+^ channels with kinetics at delta time scale^48^. At the circuit level, aside from midbrain dopaminergic inputs^45^, ChIs receive GABAergic inputs from SPNs^49^ and other local interneurons^50^, and strengthening of these synapses may also contribute to the augmentation of the delayed inhibition.

### ChI activity and ACh content

As 80–90% of cholinergic terminals lack classical synapses, ACh is known to exert both direct synaptic actions and volumetric transmission^51^. The sensory-evoked ChI responses we observed occur over a rapid time scale of ten to hundreds of milliseconds. Bulk striatal ACh concentration changes following sensory cues, as measured with GRAB-ACh sensors, are on a much slower time scale of several seconds^21,52^. Sensory-evoked immediate ChI spiking is expected to contribute to transient extracellular ACh increase, particularly locally at synapses^53^, which may be different from bulk ACh release that activate extra-synaptic receptors and trigger receptor-mediated feedback^54,55^. Thus, the immediate ChI excitation observed here may be functional distinct from the bulk ACh dynamics reported via GRAB-ACh sensors^56^.

### ACh-DA interactions in reward and aversive learning

Aversive stimuli have been found to suppress dopamine^1,2,10,11^. However, neutral-predicting stimuli can also have this effect, suggesting that dopamine neurons are insensitive to the aversiveness of stimuli, but may be suppressed by any stimulus that provides evidence against reward^2,10^. Our findings suggest that ChI excitation signals aversion that drives avoidance behavior, complementing prior work showing that DA suppression is not specific to aversiveness^2,10^. This ChI-dependent mechanism may help interpret some dopamine-related observations, including better learning from negative than positive outcomes^57^ and paradoxical kinesia in Parkinson’s disease patients with extensive DA loss^12-14^.

## Acknowledgements

We thank members of Han Lab for providing invaluable help throughout the study, Dr. Yangyang Wang and Dr. Cara Ravasio for their contributions to the data analysis scripts. X.H. acknowledges funding from NIH (1R01NS115797, 1RF1NS129520, 1R01MH122971), and NSF 1955981-CIF. E.S. acknowledges funding from NIH (T32-GM008764, F31 NS143166-01). A.L. acknowledges support from BU UROP.

The authors used an AI-assisted language tool to support editing and presentation of the manuscript text. All scientific content, interpretation, and final wording were reviewed and approved by the authors.

## Author Contributions

Y.Z., C.F. and X.H. designed the study. Y.Z. conducted the experiments, collected and analyzed the data. H.T., J.X., A.L. and A.M. assisted with data analysis. E.S. assisted with mouse surgeries. Y.Z. and X.H. prepared the manuscript, and all authors provided comments to the manuscript. X.H. supervised the study and oversaw all aspects of the project.

## Methods

### Mouse surgical preparation

All animal experiments were performed in accordance with the National Institute of Health Guide for Laboratory Animals and approved by the Boston University Institutional Animal Care and Use Committee (IACUC). Enrichment in the form of running wheels was provided to single-housed mice post-surgery. Animal facilities were maintained at around 70 °F and 50% humidity and were kept on a 12 h light/dark cycle. The study included both female and male ChAT-cre mice (*Tg(Chat-cre)GM24Gsat*, MMRRC:017269-UCD; *n* = 14, 7 females and 7 males), 8–16 weeks at the start of the experiments. 5 mice were injected with FLEX-SomArchon-BFP, 2 mice were injected with FLEX-SomArchon-GFP^34^, 7 mice were injected with DIO-ElectraOFF^34,35^.

Mice were anesthetized with 1.5–2% isoflurane and mounted on a stereotaxic frame (David Kopf Model 940). Mice were implanted with a sterilized custom imaging window with an attached infusion guide cannula. The imaging window was assembled prior to surgery by fitting a circular coverslip (size #0, outer diameter 3 mm; 64-0726 (CS-3R0); Warner Instruments) to a stainless-steel imaging cannula (outer diameter 3.17 mm, inner diameter 2.36 mm, height 2.0 mm; B004TUE45E, AmazonSupply) using ultraviolet-curable optical adhesive (Norland Optical Adhesive 60; P/N 6001; Norland Products Inc.). A guide cannula (8 mm, 26G; C315GS-5/SP; Plastics One Inc.) was soldered to the imaging cannula at 45° relative to the base of the imaging window using silver-bearing rosin-core solder (6400013; RadioShack). A craniotomy (∼3 mm diameter) was made over the right hemisphere striatum (AP +0.5 mm, ML +1.8 mm relative to the Bregma, DV -2.0 mm from pia). The overlying cortical tissue and corpus callosum were aspirated away to expose the dorsal striatum. Dental cement (C&B-Metabond® adhesive luting cement system; Parkell) was used to secure the imaging window to the skull and to mount a custom aluminum headplate posterior to the imaging window. For the 5 mice infused with FLEX-SomArchon-BFP, they additionally received 0.6μL of AAV-hSyn-Axon-jGCaMP8m into the substantia nigra (AP -3.2 mm, ML +1.25 mm, DV: -4.30 mm) during the same surgery. Due to photobleaching, GCaMP8m data was excluded from the study.

Ten days after surgery, the mice were infused through the guide cannula with viral vectors encoding SomArchon or ElectraOFF, including 1.5 μL rAAV9/CAG-FLEX-SomArchon-mTagBFP2 (titer 9.74x10^12^ GC/mL; VIGENE), 1.2 μL of rAAV8/CAG-FLEX-SomArchon-eGFP (titer 1.1x10^13^ GC/mL; Addgene) or 1.5 µL of rAAV9/CAG-DIO-ElectraOFF (titer >10^13^ GC/mL; Shanghai Sunbio Medical Biotechnology). The AAV particles were infused via a 10 μL NanoFil syringe (World Precision Instruments, LLC) coupled with an infuser (33G; C315IS-4/SPC; Plastics One, Inc.) at 100 nL/min, controlled by an infusion pump (UltraMicroPump3-4; World Precision Instruments, LLC). The infuser was left in place for 10 min to allow for virus diffusion before removal. The infusion guide was sealed by a single dummy cannula (C315DCS-5/SPC; Plastics One, Inc.) for sterilization.

### Behavior task

Mice were habituated 10–15 days after viral infusion for at least 15 min on 3–4 days per week. Mice were placed in the experimental room for 1 hour under the same room light and ambient noise as it would be for recording, followed by a habituation on the treadmill for at least 30 min prior to behavior tasks. Following habituation, mice were trained on either a delay eye-blink conditioning task or a delay water-reward conditioning task. Eye-blinking and tongue-licking behaviors were collected with a Flea USB 3.0 camera (FL3-U3-13S2C-CS; FLIR) and the Point Grey FlyCapture2 software (Ver 2.13.3) at 120 Hz. An 850-nm infrared LED illuminator (6-LED, 90° wide-angle; Univivi) was positioned ∼0.2–0.5 m from the mouse to illuminate the face and eye. Acquired mice behavior videos were analyzed offline using DeepLabCut (Ver 2.3.8)^37^. Behavioral events, recording cameras, and sensor excitation illumination were controlled by custom MATLAB scripts through a DAQ system (BNC-2110 and PCIe-6323; National Instruments). The sensor excitation illuminations were delivered from -2.01 ms before CS onset to +2.005 ms after CS onset for CS trials.

In total, 10 mice performed the aversive eye-blink conditioning task (4 males, 6 females) and 10 mice performed the water-reward conditioning task (6 males, 4 females), 6 of these mice (2 males, 4 females) completed both tasks. To counterbalance potential order effects, a subset of mice received both tasks in different sequences. 3 mice received eye-blink conditioning first followed by water-reward conditioning, whereas other 3 mice experienced the reverse order.

### Aversive conditioning task

The aversive conditioning task was conducted in darkness, with dim blue and red LED lights positioned in front of the mouse to mask the visual effects of excitation light used for GCaMP/ElectraOFF and SomArchon imaging. A continuous white noise (69–71 dB SPL) was presented as ambient background sound. A brief click sound occurred 1.1 s prior to the conditioned stimulus (CS), serving as a trial-start-cue. Each aversive CS trial consisted of a 1.1-s aCS+ sound at 74 dB SPL delivered from the left speaker, followed by a 100-ms air puff (unconditioned stimulus, US) to the left eye at 5–7 psi, or a 1.1-s aCS- sound delivered from the right speaker with no subsequent air puff. Free puff trials consisted of an air puff without any preceding trial-start-cue or CS, with sensor excitation illumination turned on 7–17 s before the puff delivery. Two distinct sounds were randomly assigned as aCS+ or aCS- across mice: a 2.5 kHz square wave with a 25% duty cycle, and a 5 kHz square wave with a 50% duty cycle presented in 25-cycle bursts at a rate of 100 bursts per second, generated by dedicated function generators (33210A; Agilent Technologies). The air puff was controlled by a solenoid valve (Clippard EV-2; Cincinnati) and delivered through a 0.5-mm cannula positioned 0.5–1 cm from the left eye. Trials were pseudo-randomized in blocks of 10 consisting of either 50% aCS+ and 50% aCS- trials, or 40% aCS+ trials, 40% aCS- trials, and 20% free puff trials, with a random inter-trial interval ranging from 27 to 37 s.

To quantify the eyelid movement during aversive conditioning task, we first used DeepLabCut to automatically select 20 representative frames per training video via the built-in k-means clustering algorithm and manually selected ∼15 frames spanning a complete blink. We manually annotated eight landmarks outlining the eyelid aperture (palpebral fissure) on these representative frames as ground truth frames. We then trained a deep neural network using ground truth frames from 2 videos per mouse across 10 recorded mice. Ground truth frames from the 20 videos were iteratively improved by refining the eyelid positions on frames that were identified as outliers by DeepLabCut during the training process. All recorded eye videos were processed using the trained network to detect eyelid landmarks. The eyelid movement was quantified as the frame-by-frame polygonal area enclosed by these landmarks. Each trial was aligned to aCS onset (or 1 s prior to air puff onset in free puff trials) and cropped from -2 s to +2 s relative to aCS onset to match the analysis window used for voltage recordings.

For each session, the natural eye area was defined as the mean eye area during the 10 s preceding session start. Trials were excluded from further analysis if the initial eye area mean (-2.5 to -2.0 s relative to aCS onset) was <40% of the natural eye area. For each remaining trial, eye area was normalized using the average measured during -2.0 to -1.5 s before aCS onset to obtain percentage eye closure, then baseline-subtracted using the mean value from -100 to 0 ms before aCS onset for subsequent behavioral analyses. For visualization of averaged eye traces, the signal was inverted so that larger values indicate greater eye closure.

To classify eye blinking during the aCS period, we applied an unsupervised clustering approach to z-normalized eyelid movement traces. Each trial’s eye trace was smoothed using a centered moving average (window size = 100 ms), then z-scored relative to its pre-aCS period (0.5 s pre-aCS to aCS onset). We then extracted a set of temporal features from each trial, including summary statistics (e.g., mean, variance, skewness), extrema, timing of peak derivatives, and signal integrals before and after the aCS. These features were standardized and reduced in dimensionality using principal component analysis (PCA), retaining the top 15 components that explained the majority of the variance. The PCA scores were then clustered using agglomerative hierarchical clustering (Euclidean distance, Ward linkage) into 32 discrete categories. Each cluster contained trials with similar temporal dynamics in eye responses, and cluster-averaged traces were visualized to examine response motifs.

In parallel, eye traces were manually reviewed by an independent observer to verify that conditioned response (‘blinks’) identified by the unsupervised procedure matched visually determined blinks. Unmatched trials were re-entered into the unsupervised classification. After two rounds, 85% of all trials showed concordance between the automated and manual classifications. Trials with a blink occurring after aCS onset and before US onset were classified as conditioned response trials (CR+), whereas trials without a detectable blink during the same period were classified as non-conditioned response (CR-).

CR onset time was defined as the first detected eye closure onset during the aCS period. The eye trace during the 1-s aCS was smoothed using a 25-ms moving window and differentiated. CR onset was taken as the time of the first prominent slope peak, identified by using `findpeaks` (MATLAB, R2023b; MathWorks) applied to the derivative with a minimum peak height of 1.5× the standard deviation of the derivative.

### Reward conditioning task

To motivate the mice, animals had one-hour access to water per day for more than 3 days before reward conditioning task. Their body weights were monitored during water restriction. The reward conditioning task was conducted in the same environment as the aversive task. A click sound same as that used in the aversive task was presented 1.1 s before the reward CSs, serving as a trial-start-cue. A white LED was illuminated from trial-start-cue to rCS onset to distinguish the two tasks. Each rCS trial consisted of a 1.0-s rCS+ sound at 74 dB SPL delivered from the left speaker, followed by a 5 µL water reward (US), or a 1.0-s rCS- sound from the right speaker with no reward. Free reward trials consisted of the reward cue and water reward without any preceding trial-start-cue, light, or CS, and with sensor excitation illumination turned on 5–11 s before reward delivery. Two distinct sounds were randomly assigned as rCS+ or rCS- across mice, including a 5 kHz square wave with a 25% duty cycle, and a 10 kHz square wave with a 50% duty cycle delivered in 50-cycle bursts at a rate of 100 bursts per second. The water reward was controlled by a solenoid valve (LHDA1233215H B; The Lee Company) and delivered simultaneously with a 100 ms reward cue (a 250 Hz square wave with 50% duty cycle, 70 dB SPL) through a ∼1.5 mm diameter tube positioned 0.3–0.5 cm from the mouth. Trials were pseudo-randomized in blocks of 10 consisting of 40% rCS+ trials, 40% rCS- trials, and 20% free reward trials, with a randomly selected inter-trial interval ranging from 6 to 12 s.

To detect the licking behavior during the reward conditioning task, we used DeepLabCut to automatically select 20 representative frames per training video. In addition, for each video we manually identified three complete lick cycles and extracted ∼20 frames spanning each cycle, yielding ∼60 manually curated frames per video. We manually annotated eight landmarks outlining the licking tongue for each representative frames as ground truth frames. The deep neural network was trained using ground truth frames from 2 videos per mouse across 10 recorded mice. All recorded lick videos were processed using the trained network to detect tongue landmarks. We calculated the tongue area as the polygonal area enclosed by the eight landmarks and summarized per-frame confidence as the mean likelihood across landmarks.

To identify licking events, we first identified high-confidence plateaus in likelihood by thresholding at 0.3 and requiring a duration of 25–125 consecutive milliseconds. For each plateau, we computed the maximum tongue area. An automatic area threshold was defaulted to the 15th percentile of the plateau maxima. Plateaus with maximum area exceeded this threshold were classified as licks.

Lick trace was quantified for each frame as the tongue area multiplied by the mean likelihood across the eight tongue landmarks. Frames with a mean landmark likelihood <0.3 were set to zero. For session-wise analysis, The resulting lick trace was normalized to the mean of the top 15% of maximal values detected within 1 s after US delivered.

A conditioned response trial (CR+) in the reward task was defined as at least two consecutive licks occurring after rCS onset and before US (water delivery). Trial with zero licks during the same time window was classified as CR-. In CR+ trials, the first frame of earliest accepted lick plateau occurring during rCS was defined as CR onset time.

### Behavioral metrics

Behavioral performance was evaluated by hit rate and d-prime. Hit rate was calculated as CS+ trials with CR+ divided by all CS+ trials. d-prime was calculated as

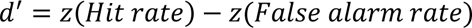

where rates of 0 or 1 were corrected using 0.5/(N+1) prior to z-scoring (N is the number of trials in that condition). Mice typically learned these tasks in ∼1week, reaching a behavioral threshold defined as having a hit rate >50% and a d-prime > 0.7 for two consecutive days.

### Optogenetic silencing of ChI during conditioning tasks

For optogenetic manipulation experiments, ChI activity was transiently suppressed using the light-driven silencer ArchT^58^ during defined time windows of the conditioned stimulus (CS). Mice were infused with 1.2 μL of rAAV9/CAG-Flex-ArchT-GFP (titer 4.4x10^12^; Gene Therapy Center Vector Core) via the infusion guide cannula 10 days after window implantation surgery. Habituation and conditioning tasks commenced 10 days after viral infusion.

Six mice (3 males, 3 females) were trained in the aversive and reward conditioning tasks similar to those described above until the post-learning stage prior to optogenetic manipulation. In optogenetic silencing experiment, the task consisted only of CS+ and CS- trials (50% each, pseudo-randomized in blocks of 10), and free trials were not included. To counterbalance potential order effects, 3 mice received eye-blink conditioning first followed by water-reward conditioning, while other 3 mice experienced the reverse order. One mouse that failed to reach the behavioral threshold after 15 training days was excluded from the experiment. Behavioral video acquisition and analysis were performed using the same procedures described above for the aversive and reward conditioning tasks.

After animals reached the post-learning stage, optogenetic silencing experiments were conducted over 7 consecutive days, including immediate ChI silencing for 4 days followed by delayed silencing for 3 days. Silencing was achieved using ArchT activated by a 565 nm LED (M565L3; Thorlabs). Light was focused via a microscope objective (CFI Apo NIR 40×/0.8NA W; Nikon) and delivered through the recording window at an intensity of ∼100 mW/mm^2^ at the dorsal striatum. Silencing was applied either during immediate window (0–300 ms after CS onset) or late window (300–600 ms after CS onset) in separate sessions. Within each session, ArchT-ON and ArchT-OFF trials were interleaved (50% each, pseudo-randomized in blocks of 10, independent of CS identity), allowing within-session comparisons of behavioral.

### In vivo imaging

*In-vivo* imaging in head-fixed behaving mice was performed at least 2 weeks after virus infusion for sufficient sensors expression. Voltage and calcium imaging were performed using a custom dual-channel wide-field fluorescence microscope. A 470 nm LED (M470L4; Thorlabs) for Axon-GCaMP or ElectraOFF excitation and a 637 nm laser (Necsel Red 63×; Ushio America) for SomArchon excitation were combined using a 505 nm long-pass filter (DMLP505R; Thorlabs) and focused onto the sample via a microscope objective (CFI Apo NIR 40×/0.8NA W; Nikon). The intensity of 637-nm laser excitation on the imaging field was kept ∼10 W/mm^2^, and the intensity of the 470-nm LED excitation for GCaMP8m was kept ∼50 mW/mm^2^. The intensity of the 470-nm LED excitation for ElectraOFF was kept ∼150 mW/mm^2^.

The generated fluorescence signals were collected by the same objective and separated from the excitation sources using a multi-band filter set (LF405/488/532/635-B; Semrock). To enable simultaneous two-color imaging, GCaMP and SomArchon fluorescence were split into two separate paths using a multi-band dichromatic mirror (ZT405/514/635rpc; Chroma Technology Corp.). GCaMP/ElectraOFF fluorescence signal was filtered with a 525 ± 22.5 nm bandpass emission filter (FF01-525/45; Semrock) and recorded by a sCMOS camera (C13440-20CU; Hamamatsu Photonics). SomArchon fluorescence signal was filtered with a 700 ± 37.5 nm bandpass emission filter (ET700/75m; Chroma Technology Corp.) and recorded by second sCMOS camera (C15440-20UP; Hamamatsu Photonics).

To measure membrane voltage from individual striatal neurons during conditioning learning tasks, we triggered the SomArchon camera via TTL pulses generated by MATLAB via a DAQ. Once triggered, HCImage Live (version 4.8.0; Hamamatsu Photonics) collected 3 or 4 s of data. HCImage Live was set to capture an image frame of 144×144 or 144×288 pixels with 2×2 binning, corresponding to 46×46 µm^2^ or 46×92 µm^2^, with an exposure time of 2 ms for SomArchon. When the SomArchon camera was first triggered, it generated a TTL pulse and sent it to the GFP camera and HCImage Live, and continuously collected same length of data in a field of view (FOV) at 1024×1024 pixels with 2×2 binning, corresponding to 328×328 µm^2^, with an exposure time set at 50 ms for GCaMP8m. For ElectraOFF, HCImage Live was set to capture an image frame of 128×256/512/1024 pixels with 2×2 binning, corresponding to 41×82/164/328 µm^2^, with an exposure time of 1.5 ms. The actual frame capture from the SomArchon and GFP cameras, delivery of trial-start-cue/CS/US, were simultaneously monitored as TTL pulses with an Open Ephys acquisition board (SKU: OEPS-9030; Open Ephys) at a rate of 30 kHz to ensure proper alignment during further analysis offline. Acquired imaging data of every trial were stored in dcimg format for off-line analysis using MATLAB.

### Motion correction and trace extraction method

All fluorescence video analyses were performed offline using custom scripts. Image stacks were subjected to motion correction using a pairwise rigid motion correction algorithm as described previously^59^. Briefly, frame-by-frame shifts were estimated by computing the maximum cross-correlation coefficient between each frame and the reference image, defined as the average of all frames within each individual trial. Regions of interest (ROIs) corresponding to individual neurons were manually delineated using the `drawPolygon` function (MATLAB) from the trial-averaged image after motion-corrected. The raw fluorescence trace of individual neuron was extracted by averaging pixel intensities within the ROI. The background trace was extracted from a separate ROI selected in the same manner from an area devoid of the cell bodies and dendrites. Frames with background trace z-scores greater than 10 or less than -10 were flagged as motion artifacts and excluded prior to further analysis.

### Background subtraction and Spike detection

To minimize the effects of photobleaching, neuronal and background traces from SomArchon were first detrended using `detrend` function (MATLAB). The background component was then estimated using robust linear regression (poly1, LAR method in MATLAB) to minimize the influence of transient outliers. ElectraOFF traces were not detrended due to their high photostability^35^. The background component was estimated using ordinary least-squares regression (`fitlm`, MATLAB) subtracted from the raw neuronal trace. For signals collected with ElectraOFF, which reports membrane depolarization as a decrease in fluorescence, traces were sign-inverted so that upward deflections correspond to depolarization. For signals collected with SomArchon, which reports depolarization as an increase in fluorescence, traces were not inverted. Finally, each trial’s refined trace was median-centered by subtracting its median.

Spikes were detected from the median-centered refine Vm trace. We first smoothed the Vm trace using moving-average (150 ms window) and Savitzky-Golay smoothing (2nd order, 150 ms window) separately, then averaged the two resulting traces to obtain a composite baseline. The composite baseline was subtracted from the Vm trace to extract the high-frequency Vm component (Vm_high_). To estimate noise while minimizing contamination by spikes, we median-centered the Vm_high_ signal and calculated the standard deviation of its negative fluctuations. Spikes were detected using MATLAB `findpeaks` function as peaks exceeding 4.5x noise standard deviations for SomArchon recordings and 5.5x noise SD for ElectraOFF recordings. Spike amplitude was calculated using the Vm trace as the difference between the spike peak and the minimum fluorescence value within the ∼7 ms (3 frames for SomArchon and 5 frames for ElectraOFF) preceding the spike peak. Spikes with an amplitude lower than 3x SD were considered as misdetections and excluded for further analysis. Firing rate (FR) was computed from the remaining spikes and smoothed using a 6-ms moving window.

### Quantification of neuronal Vm and FR responses

Vm and FR traces recorded from SomArchon expressing neurons were interpolated to a frame rate of 666 Hz to match that of the ElectraOFF expressing neurons. For population analysis, each trial’s Vm trace was z-scored across the entire trial. Event-evoked Vm and FR changes were quantified by subtracting the pre-event baseline from -100 to 0 ms relative to event onset. Neurons were identified across sessions based on cell morphology and surrounding vasculature structures. When a neuron was recorded in multiple sessions within the same task and learning stage, the neuronal Vm and FR traces were computed by averaging the trial-wise means across sessions. For neurons recorded once in the task and learning stage, the neuronal Vm and FR traces were computed by averaging across trials.

Because SomArchon recordings in the aversive task did not include the trial-start-cue period, comparisons of trial-start-cue responses between aversive and reward tasks were restricted to ElectraOFF-expressing neurons. Due to photobleaching, free puff trials were not acquired with SomArchon. Free-trial analyses were therefore restricted to ElectraOFF-expressing neurons for consistency across aversive and reward conditions.

Response latency to task-relevant events was determined from neuronal Vm and FR traces as the first sustained suprathreshold response relative to baseline variability. For each neuron, the baseline mean and standard deviation were calculated from the -100 to 0 ms pre-event window, and the trace was converted to a z-score relative to this baseline. Response detection was restricted to a post-event window within 30 ms after event onset. Response onset was defined as the first time point at which the z-scored trace exceeded 2.5 standard deviations above baseline for at least 5 ms. Latency was defined as the interval between event onset and the detected response onset and is reported in **Table S1**.

Neurons were classified as significantly modulated by task-relevant event if the mean Vm (or FR) during the immediate window (35–100 ms after event onset) differed from the pre-event baseline (-100 to 0 ms relative to event onset) across trials (Wilcoxon signed-rank test, *p* < 0.05). The fraction of modulated neurons is reported in **Table S2**. Neurons were classified as discriminating between the two CSs if the mean evoked Vm (or FR) during the immediate response window was significantly different for CS+ and CS- trials (Mann-Whitney U tests, *p* < 0.05). The fraction of discriminating neurons is reported in **Table S3** and the 10 ChI recorded across aversive and reward task in late learning are reported in **Table S4**.

## Supplement Information

**Fig. S1.**
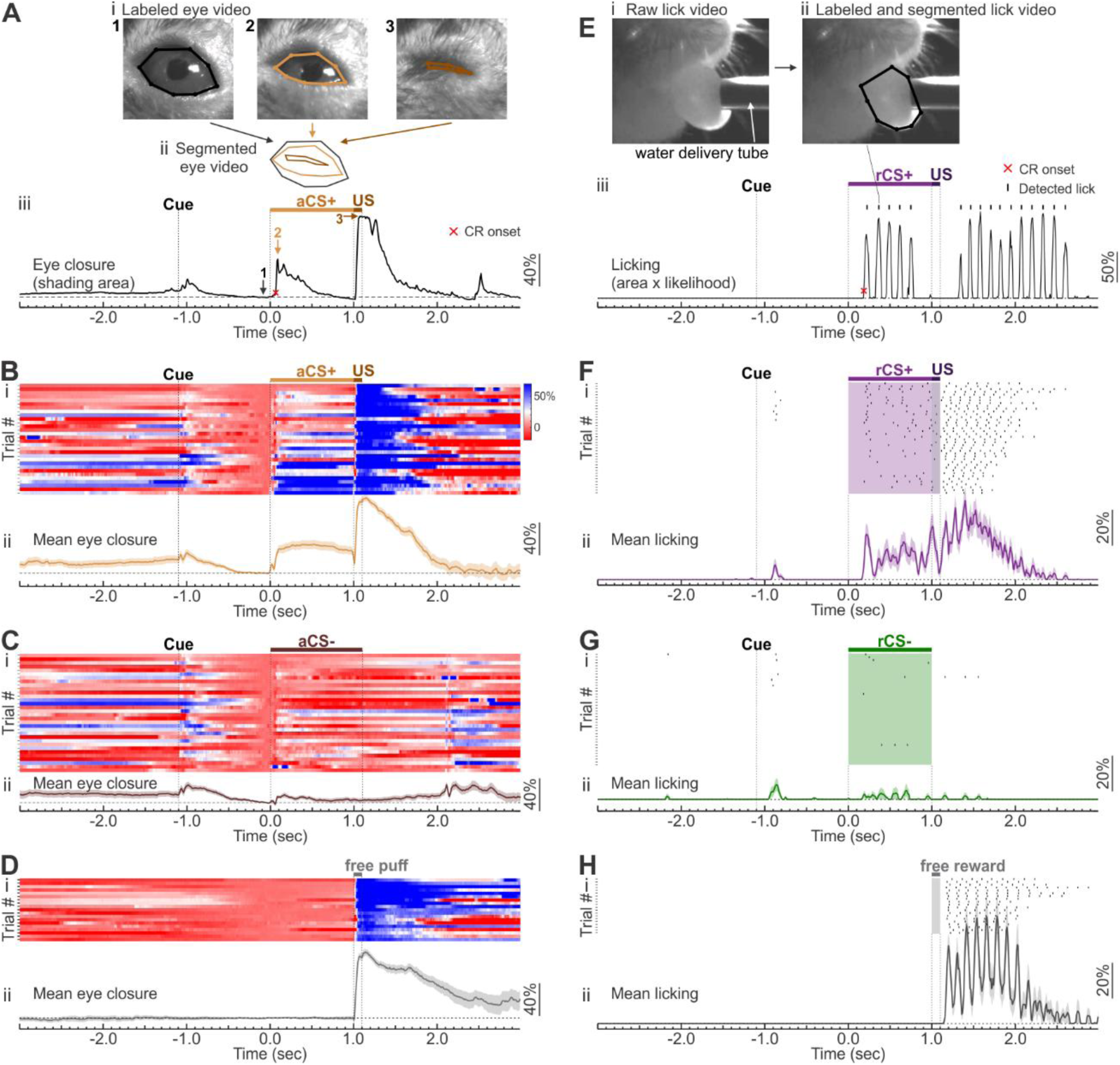
Quantification of animal behavior. (**A**) Example aCS+ trial showing eye monitoring and eye closure quantification. (**Ai**) Representative frames with labelled eyes size aligned to the CS onset. Frame 1 is before CS onset with eye open, frame 2 is during aCS+ showing partial eye closure, frame 3 is during US showing stronger closure. (**Aii**) Segmented eye area used to calculate eye closure. (**Aiii**) Eye closure (%, normalized and baseline-subtracted) from the example trial. Vertical dotted line marks trial-start-cue onset. The orange bar denotes the aCS+ period, followed by US delivery. Dashed line is at 0, corresponding to the mean value during -100 ms to 0 ms pre-CS baseline. The numbered arrows correspond to the three example frames above. Red cross indicates CR onset. (**Bi**) Heatmap of eye closure (%) across aCS+ trials in late aversive learning (*n* = 30 trials). (**Bii**) Mean eye closure across the above trials. Solid lines indicate the mean and shaded areas represent SEM. (**C**) Similar to (B) but for the aCS- trials from the same session (*n* = 32 trials). (**D**) Similar to (B) but for the free puff trials from the same session (*n* = 19 trials). (**E**) Example rCS+ trial showing tongue monitoring and lick detection. (**Ei**) Representative raw lick frame during CS. (**Eii**) Labelled lick frame and segmented tongue area used to detect licks. (**Eiii**) Lick trace (% tongue area multiplied by the mean landmark likelihood) from the example trial with detected lick events. Vertical dotted line marks trial-start-cue onset. The purple bar denotes the rCS+ period, followed by US delivery. Dashed line is at 0. Detected lick events are shown as raster ticks above the trace. Red cross indicates CR onset. (**Fi**) Trial-by-trial raster of detected lick events during rCS+ trials in late reward learning (*n* = 40 trials). (**Fii**) Mean lick traces across the above trials. (**G**) Similar to (F) but for the rCS- trials from the same session (*n* = 40 trials). (**H**) Similar to (F) but for the free reward trials from the same session (*n* = 20 trials).

**Fig. S2.**
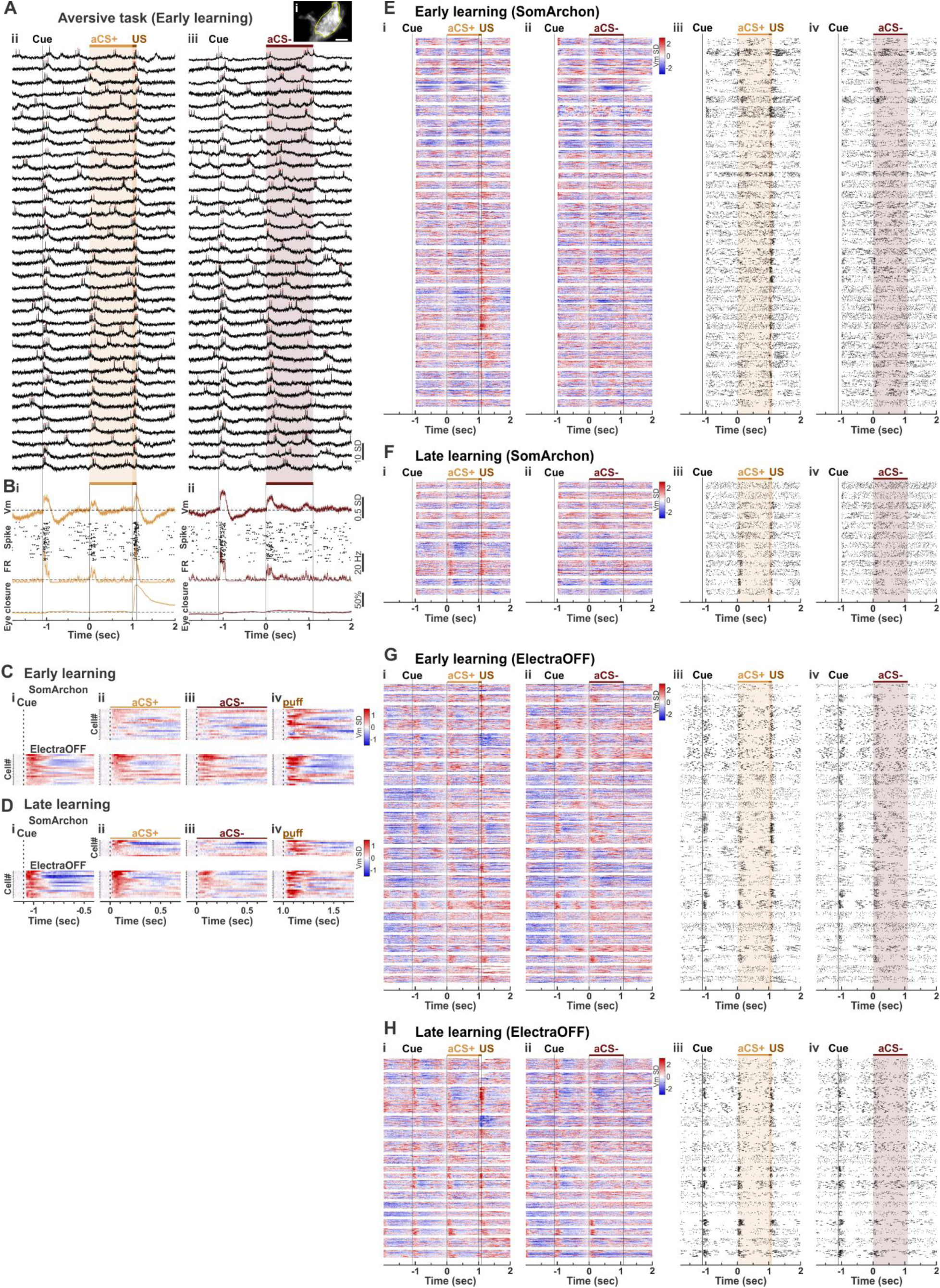
Individual ChIs responses during aversive learning. (**A, B**) An example recording session from one ElectraOff expressing ChI during the aversive task in early learning. (**Ai**) Upper right: ElectraOFF fluorescence image, averaged across all image frames during one trial. Yellow outline indicates the ROI selected as the ChI soma; scale bar, 10 μm. (**Aii-iii**) Vm traces (black) across example trials aligned to CS onset. Red dots indicate spikes. (**Aii**) Aversive CS+ trials. CS+ duration is highlighted by orange shading. US: dark brown shading. (**Aiii**) Aversive CS- trials. CS- duration is highlighted by maroon shading. Trial-start-cue is labelled as vertical dashed line. (**B**) Vm, FR overlaid with spike raster, and eye closure across all aCS+ (**Bi**, orange: *n =* 35 trials) and aCS- (**Bii,** maroon: *n =* 35 trials) trials. Vm, FR, and eye closure were baseline-subtracted using the corresponding mean during -100 ms to 0 ms pre-CS baseline for each trial. Dashed lines are at 0. Solid lines indicate the mean and shaded areas represent SEM. (**C**) Heatmaps of the evoked Vm by trial-start-cue (Ci), aCS+ (Cii), aCS- (Ciii), and US (Civ) across all neurons in early learning. SomArchon expressing neurons (*n* = 19 neurons from 5 mice) on top and ElectraOFF expressing neurons (*n* = 19 neurons from 5 mice) on bottom. Each row represents one neuron. Evoked Vm was computed as the trial-wise mean of the baseline subtracted Vm, with the baseline defined as the mean during the -100 to 0 ms pre-event period for each trial. Neurons are sorted by the amplitude of the evoked Vm by aCS+ during immediate window. (**D**) Similar to (C) but for late learning, with *n* = 10 SomArchon neurons from 4 mice and *n* = 17 ElectraOFF neurons from 4 mice. (**E**) Heatmaps of the evoked Vm responses (baseline-subtracted and z-scored, Ei-Eii) and spike raster (Eiii-Eiv) across all SomArchon neurons in early learning. Each row represents one trial, trials from the same neuron are shown in a block. Neurons are shown in recording order. Responses to the aCS+ (Ei, Eiii, orange) and aCS- (Eii, Eiv, maroon) are shown for the same neurons. (**F**) Similar to (E) but for late learning. (**G-H**) Similar to (E-F) but for all ElectraOFF neurons.

**Fig. S3.**
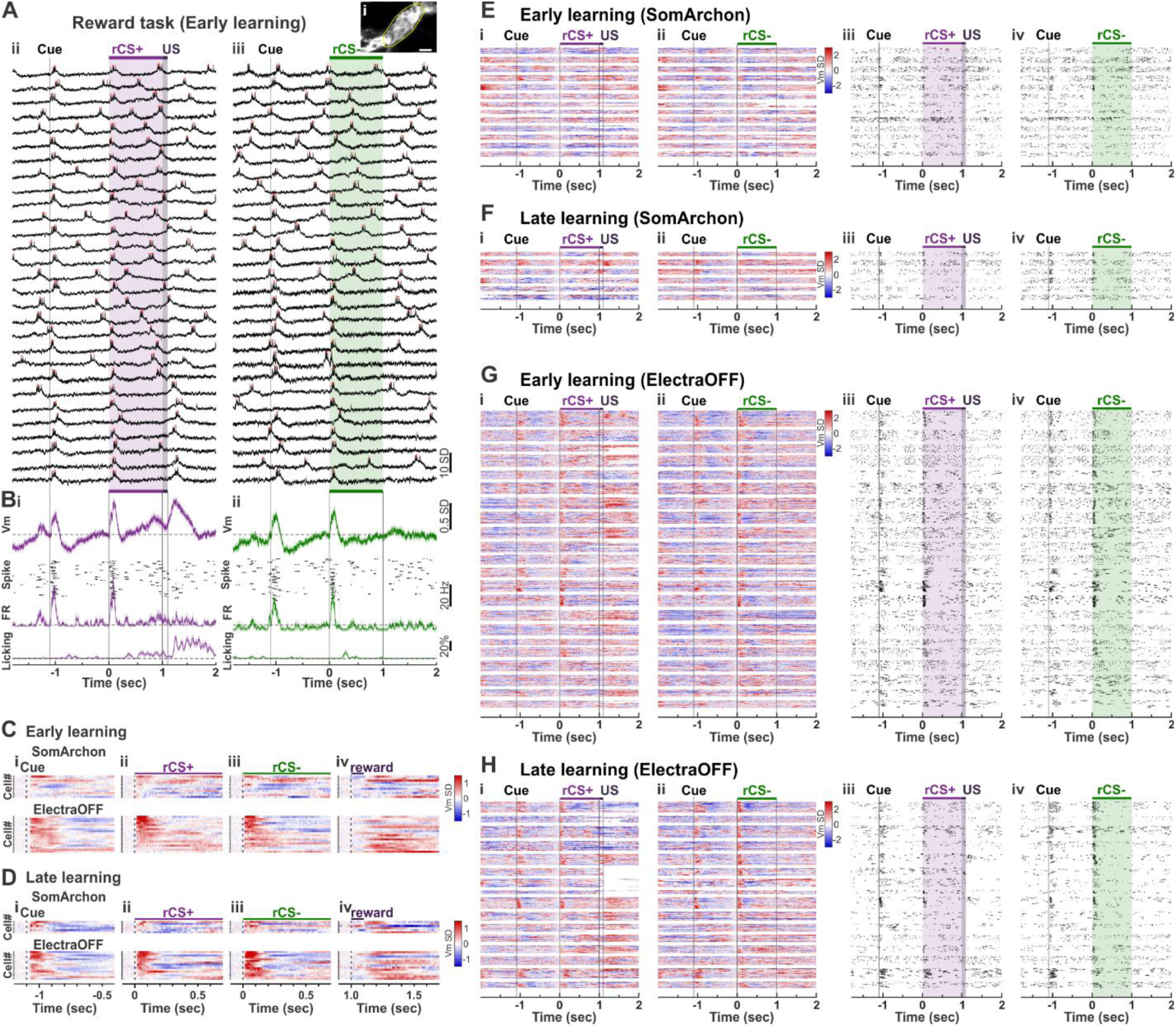
Individual ChIs responses during reward learning. (**A, B**) An example recording session from one ElectraOff expressing ChI during the reward task in early learning. (**Ai**) Upper right: ElectraOFF fluorescence image, averaged across all image frames during one trial. Yellow outline indicates the ROI selected as the ChI soma; scale bar, 10 μm. (**Aii-iii**) Vm traces (black) across example trials aligned to CS onset. Red dots indicate spikes. (**Aii**) Reward CS+ trials. CS+ duration is highlighted by purple shading. US: dark purple shading. (**Aiii**) Reward CS- trials. CS- duration is highlighted by green shading. Trial-start-cue is labelled as vertical dashed line. (**B**) Vm, FR overlaid with spike raster, and tongue licking across all rCS+ (**Bi**, purple: *n =* 29 trials) and rCS- (**Bii,** green: *n =* 29 trials) trials. Vm, FR, and tongue licking were baseline-subtracted using the corresponding mean during -100 ms to 0 ms pre-CS baseline for each trial. Dashed lines are at 0. Solid lines indicate the mean and shaded areas represent SEM. (**C**) Heatmaps of the evoked Vm by trial-start-cue (Ci), rCS+ (Cii), rCS- (Ciii), and US (Civ) across all neurons in early learning. SomArchon expressing neurons (*n* = 11 neurons from 3 mice) on top and ElectraOFF expressing neurons (*n* = 18 neurons from 7 mice) on bottom. Each row represents one neuron. Evoked Vm was computed as the trial-wise mean of the baseline subtracted Vm, with the baseline defined as the mean during the -100 to 0 ms pre-event period for each trial. Neurons are sorted by the amplitude of the evoked Vm by rCS+ during immediate window. (**D**) Similar to (C) but for late learning, with *n* = 6 SomArchon neurons from 3 mice and *n* = 16 ElectraOFF neurons from 5 mice. (**E**) Heatmaps of the evoked Vm responses (Ei-Eii) and spike raster (Eiii-Eiv) across all SomArchon neurons in early learning. Each row represents one trial, trials from the same neuron are shown in a block. Neurons are shown in recording order. Responses to the rCS+ (Ei, Eiii, purple) and rCS- (Eii, Eiv, green) are shown for the same neurons. (**F**) Similar to (E) but for late learning. (**G-H**) Similar to (E-F) but for all ElectraOFF neurons.

**Fig. S4.**
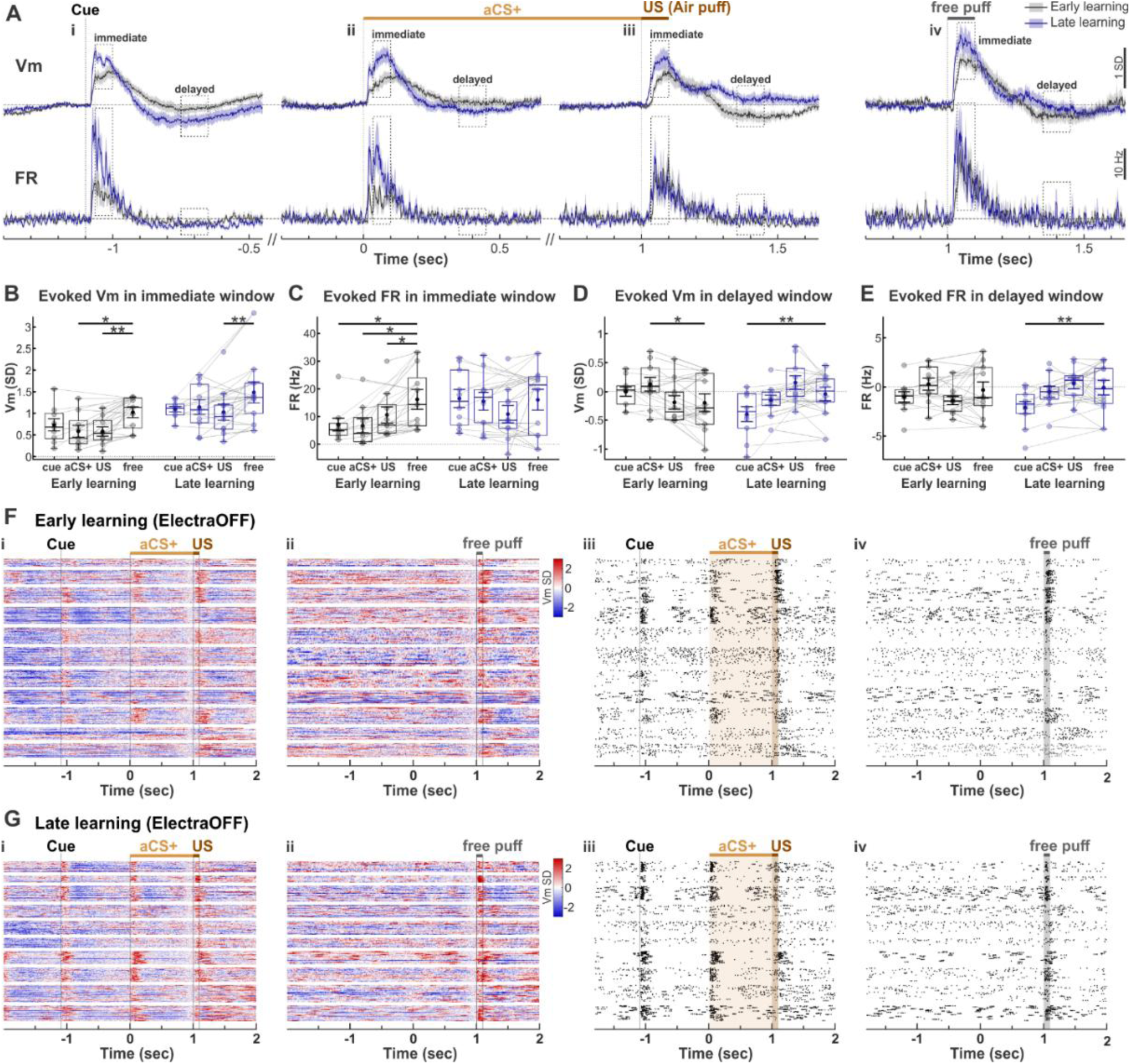
ChIs responses to free puff. (**A**) Population Vm, FR across recorded neurons aligned to trial-start-cue (Ai), aversive CS+ (Aii), US (Aiii), and free puff (Aiv) during early (grey) and late (blue) learning. Dash lines are at 0. Solid lines indicate mean and shaded areas represent SEM (early learning, *n =* 10 neurons from 4 mice; late learning, *n =* 10 neurons from 3 mice). Immediate and delayed windows are highlighted by the dashed boxes. (**B-E**) Event-evoked Vm **(B, D)** and FR (**C, E**) during the immediate (**B, C**) and delayed (**D, E**) windows for trial-start-cue, aCS+, US and free puff during early (grey) and late (blue) learning. Boxes indicate the interquartile range (IQR; 25th–75th percentiles) across neurons. Center lines are the median, and whiskers are the 1.5 × IQR. Mean ± SEM are overlaid on the box plots. Horizontal lines with asterisks indicate comparisons between different conditions and free puff using Wilcoxon signed-rank tests (see **Table S6** for statistical details). (**F**) Heatmaps of the evoked Vm responses (baseline-subtracted and z-scored, Fi-Fii) and spike raster (Fiii-Fiv) across all neurons with free puff trials in early learning. Each row represents one trial, trials from the same neuron are shown in a block. Neurons are shown in the recording order. Responses to the aCS+ (Fi, Fiii, orange), and free puff (Fii, Fiv, grey) are shown for the same neurons. (**G**) Similar to (F) but for late learning. * *p <* 0.05, ** *p <* 0.01, *** *p <* 0.001.

**Fig. S5.**
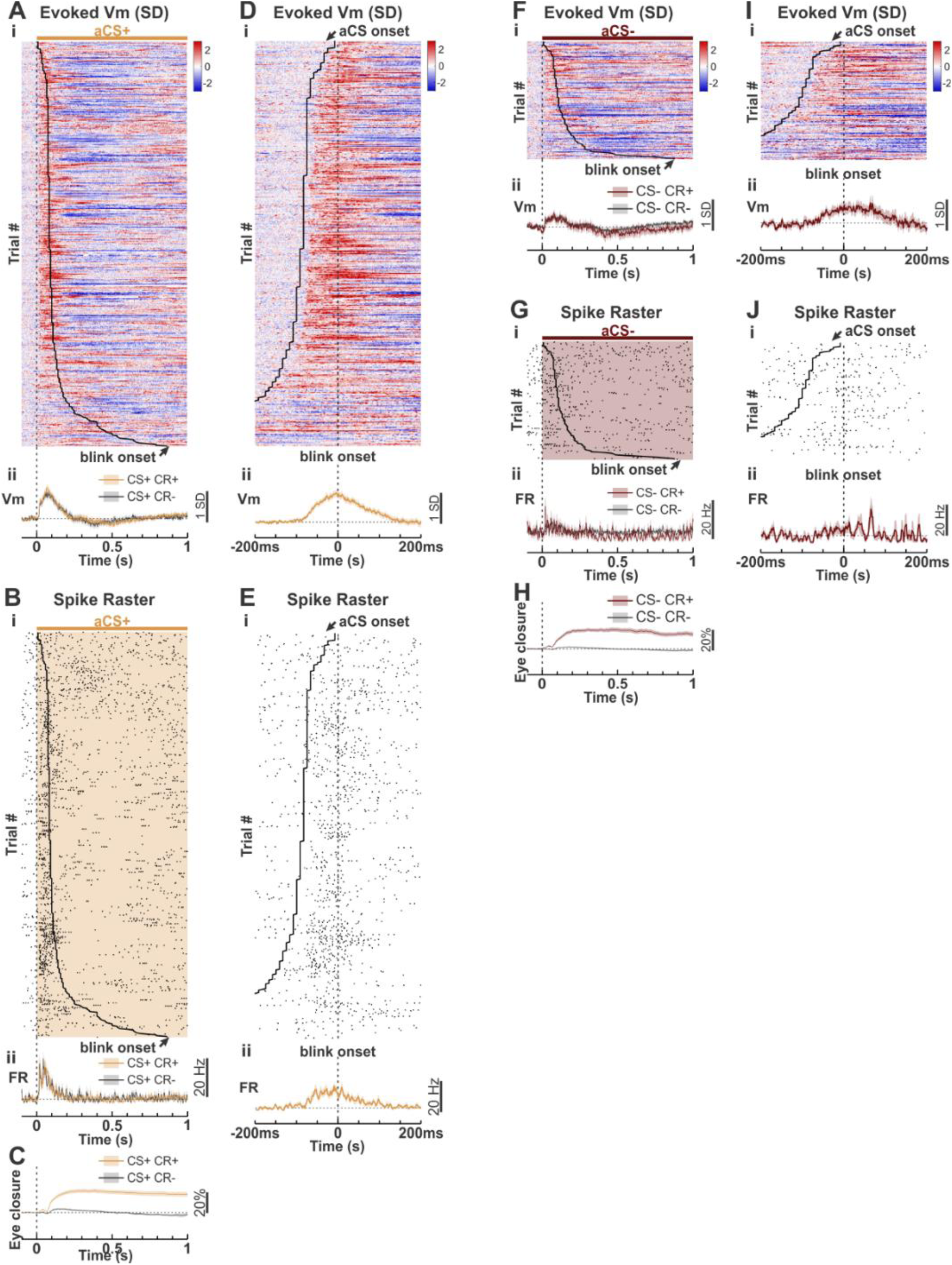
During aversive learning, ChI activation is time locked to aCS onset, but not to movement onset. (**A**) Heatmaps of evoked Vm responses (Ai) aligned to aversive CS+ onset (vertical dashed line) across 522 CS+ CR+ trials in late learning. Each row represents one trial, trials are sorted by CR onset time (black curve). (**Aii**) Vm across all CS+ CR+ (orange) and CS+CR- (grey) trials (*n* = 27 neurons from 8 mice). Dashed lines are at 0. Solid lines indicate the mean and shaded areas represent SEM. (**B**) Similar to (A) but for spike raster and FR. (**C**) Similar to (Aii) but for eye closure. (**D**) Heatmap (Di) and neuron-wise mean (Dii) of the Vm responses from the same trials as (A), but aligned to CR onset (vertical dashed line). CS onset time was labelled as black curve. (**E**) Similar to (D) but for spike raster and FR. (**F-J**) Similar to (A-E) but for 152 CS- CR+ trials (maroon) from 22 neurons, and 27 neurons for CS- CR- (grey) from 8 mice.

**Fig. S6.**
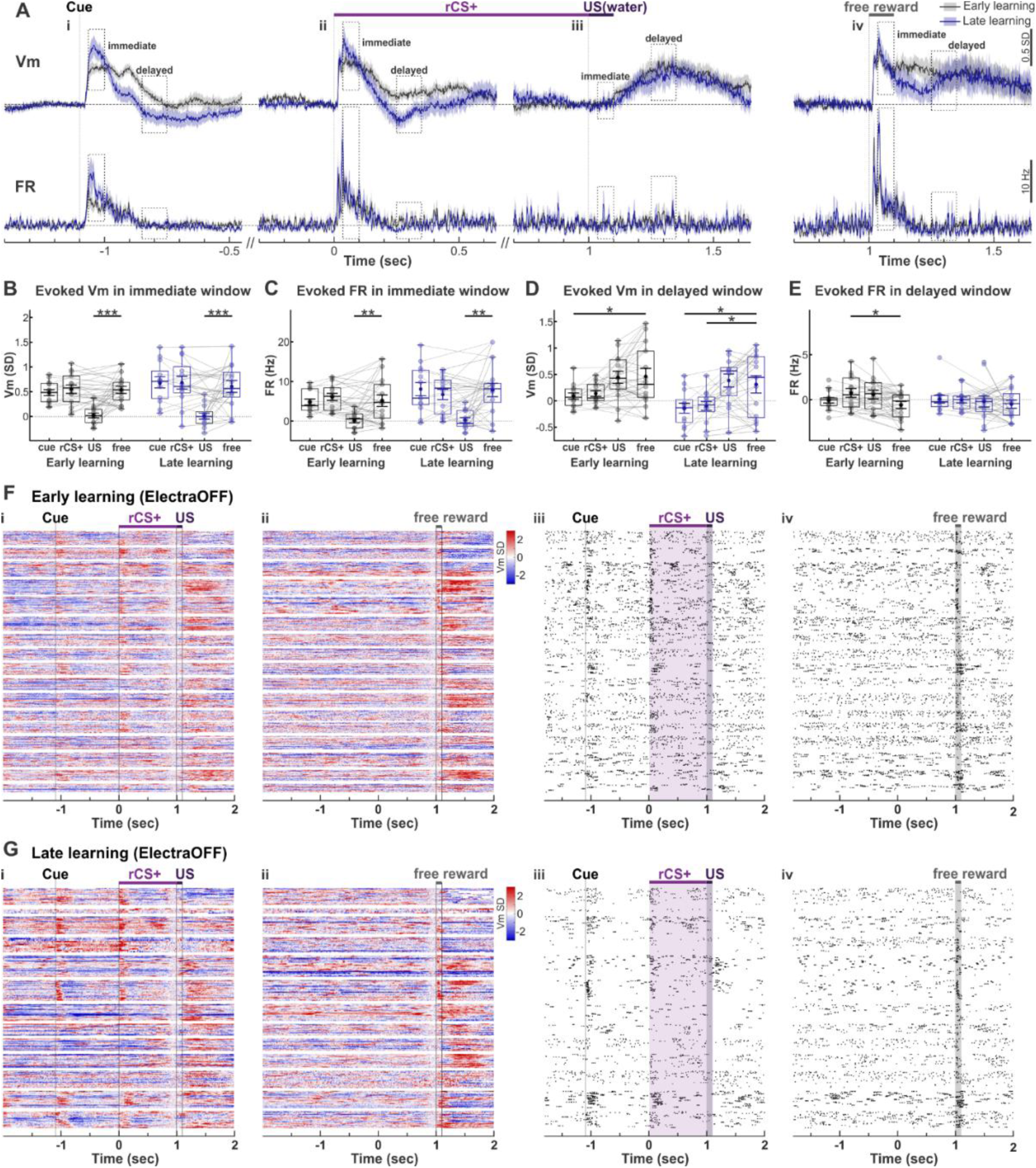
ChIs responses to free reward. (**A**) Population Vm, FR across recorded neurons aligned to trial-start-cue (Ai), reward CS+ (Aii), US (Aiii), and free reward (Aiv) during early (grey) and late (blue) learning. Dash lines are at 0. Solid lines indicate mean and shaded areas represent SEM (early learning, *n =* 14 neurons from 6 mice; late learning, *n =* 13 neurons from 5 mice). Immediate and delayed windows are highlighted by the dashed boxes. (**B-E**) Event-evoked Vm **(B, D)** and FR (**C, E**) during the immediate (**B, C**) and delayed (**D, E**) windows for trial-start-cue, rCS+, US and free reward during early (grey) and late (blue) learning. Boxes indicate the interquartile range (IQR; 25th–75th percentiles) across neurons. Center lines are the median, and whiskers are the 1.5 × IQR. Mean ± SEM are overlaid on the box plots. Horizontal lines with asterisks indicate comparisons between different conditions and free reward using Wilcoxon signed-rank tests (see **Table S7** for statistical details). (**F**) Heatmaps of the evoked Vm responses (baseline-subtracted and z-scored, Fi-Fii) and spike raster (Fiii-Fiv) across all neurons with free reward trials in early learning. Each row represents one trial, trials from the same neuron are shown in a block. Neurons are shown in the recording order. Responses to the rCS+ (Fi, Fiii, purple), and fre e reward (Fii, Fiv, grey) are shown for the same neurons. (**G**) Similar to (F) but for late learning. * *p <* 0.05, ** *p <* 0.01, *** *p <* 0.001.

**Fig. S7.**
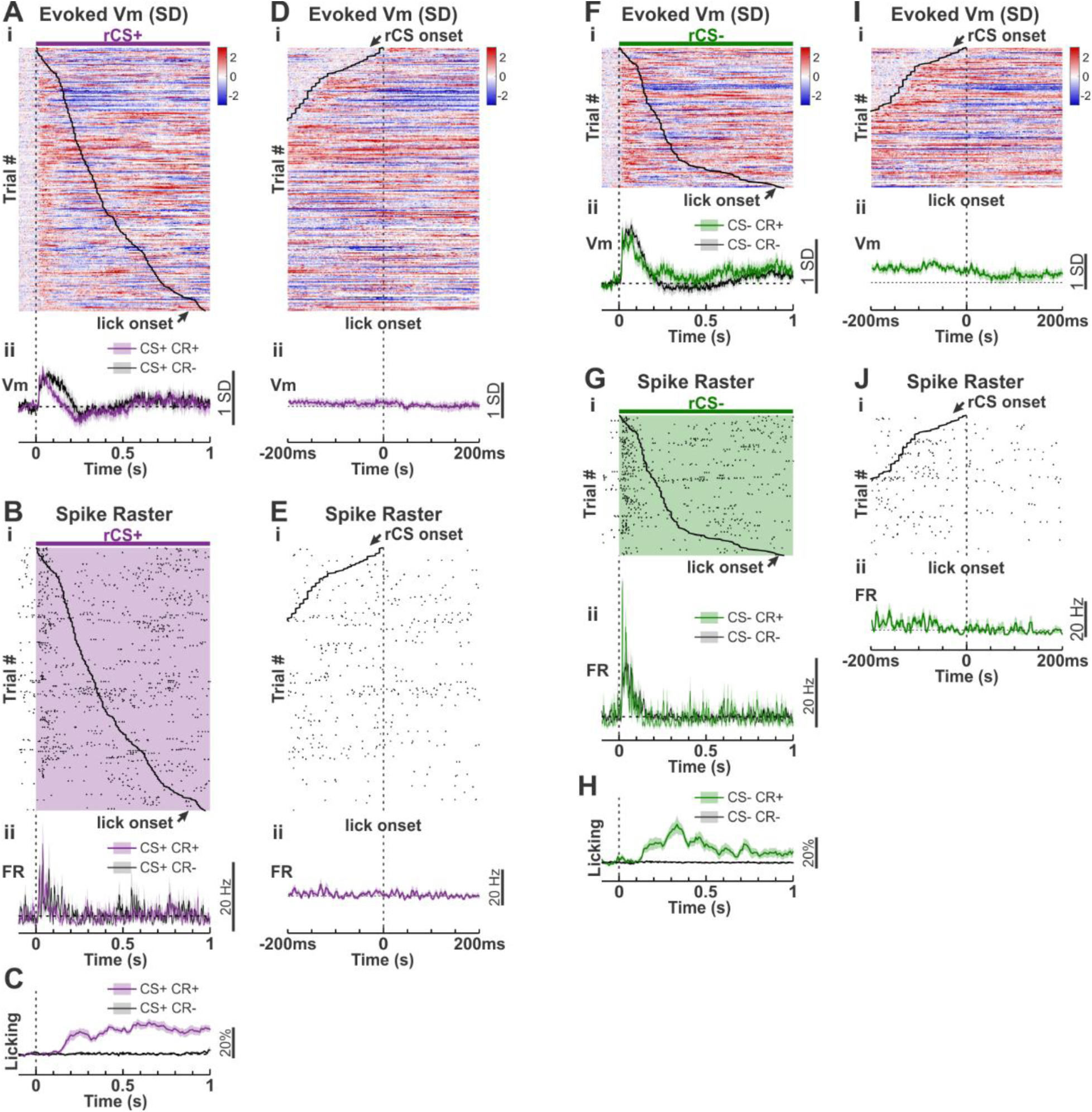
During reward learning, ChIs did not respond to lick movement, despite robust response to rCS. (**A**) Heatmaps of evoked Vm responses (Ai) aligned to reward CS+ onset (vertical dashed line) across 295 CS+ CR+ trials in late learning. Each row represents one trial, trials are sorted by CR onset time (black curve). (**Aii**) Vm across all CS+ CR+ (purple, *n* = 22 neurons from 8 mice) and CS+CR-(grey, *n* = 14 neurons from 8 mice) trials. Dashed lines are at 0. Solid lines indicate the mean and shaded areas represent SEM. (**B**) Similar to (A) but for spike raster and FR. (**C**) Similar to (Aii) but for tongue licking. (**D**) Heatmap (Di) and neuron-wise mean (Dii) of the Vm responses from the same trials as (A), but aligned to CR onset (vertical dashed line). CS onset time was labelled as black curve. (**E**) Similar to (D) but for spike raster and FR. (**F-J**) Similar to (A-E) but for 156 CS- CR+ trials (green) from 16 neurons, and 22 neurons for CS- CR- (grey) from 8 mice.

**Fig. S8.**
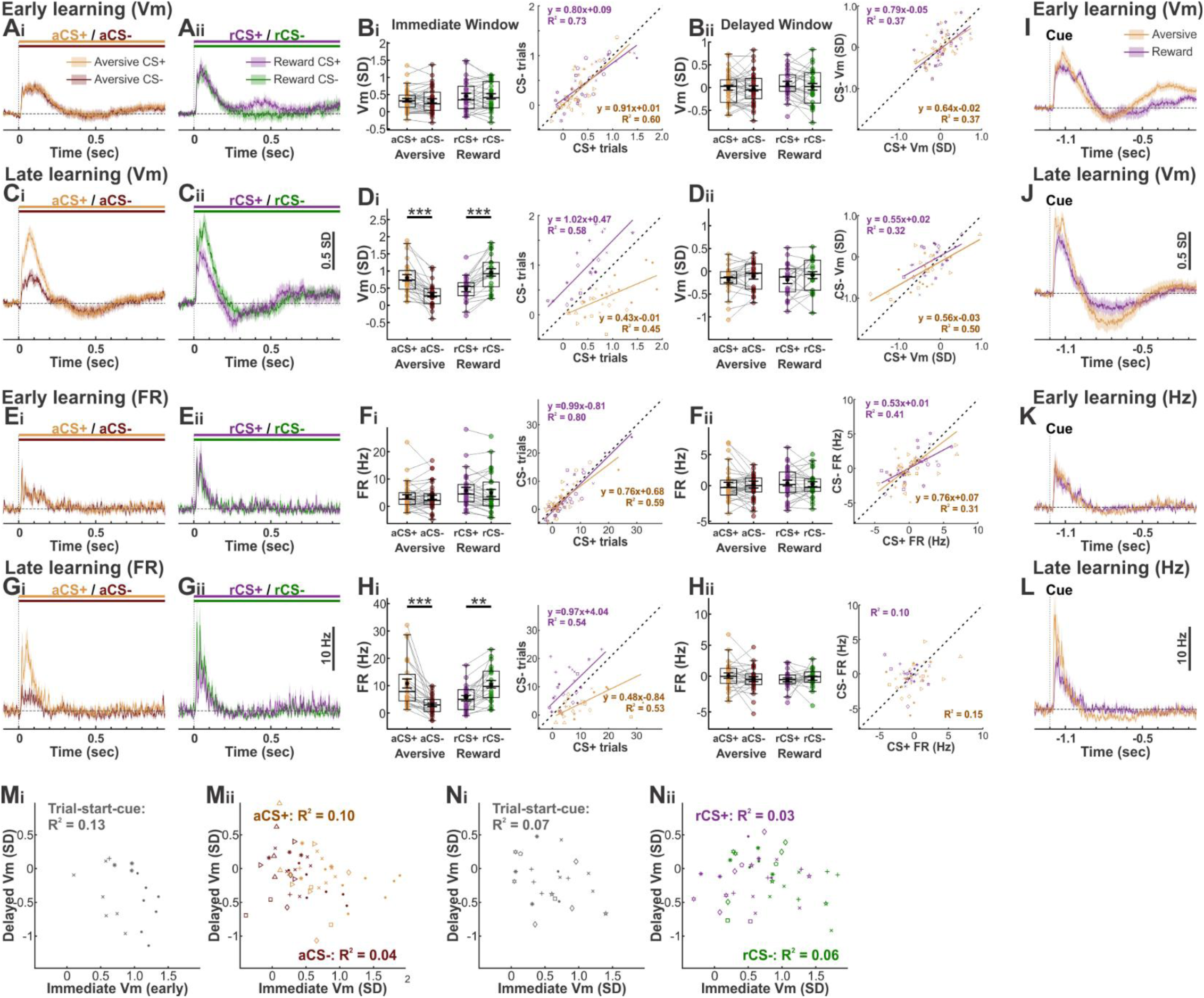
ChIs Vm and FR response to trial-start-cue and CSs in aversive and reward tasks. (**A**) Population Vm across recorded neurons aligned to CS+ and CS- in aversive (Ai, *n =* 38 neurons from 10 mice) and reward (Aii, *n =* 29 neurons from 10 mice) tasks during early learning stages. Dash lines are at 0. Solid lines indicate mean and shaded areas represent SEM. (**B**) Vm during the immediate (Bi) and delayed (Bii) window in the two tasks. Boxes indicate the interquartile range (IQR; 25th–75th percentiles) across neurons. Center lines are the median, and whiskers are the 1.5 × IQR. Mean ± SEM are overlaid on the box plots. Horizontal lines with asterisks indicate comparisons between CS+ and CS- using Wilcoxon signed-rank test (see **Table S9** for statistical details). Scatter plots of each individual neuron response to CS+ versus CS- when the mice were late learning during aversive (orange) and reward (purple) conditioning tasks. Symbols indicate different animals. Solid line indicates linear regression, with equation and R^2^ shown in panel (**C-D**) Similar to (A-B) but during late learning stage. (**E-H**) Similar to (A-D) but for FR. (**I-L**) Population Vm (I-J) and FR (K-L) across recorded neurons aligned to trial-start-cue in aversive (orange) and reward (purple) tasks during early (I, K) and late (J, L) learning (Early learning, *n =* 19 neurons from 4 mice for aversive; *n =* 18 neurons from 7 mice for reward. Late learning, *n =* 17 neurons from 4 mice for aversive; *n =* 16 neurons from 5 mice for reward). (**M-N**) Correlation of event-evoked immediate Vm depolarization and delayed Vm hyperpolarization across individual neurons in late aversive (**M**) and reward (**N**) learning, including trial-start-cue (**Mi, Ni**) and both CSs (**Mii**, orange for aversive CS+ and maroon for aversive CS-; **Nii**, purple for reward CS+ and green for reward CS-). Only R² values are shown due to weak linear fits.

**Table S1.** Summary of neuronal response latencies to task-relevant events in the two tasks. Values are mean ± SEM (ms) across neurons.

|  |  | Aversive |  |  | Reward |  |  |
| --- | --- | --- | --- | --- | --- | --- | --- |
|  |  | Cue | aCS+ | aCS- | Cue | rCS+ | rCS- |
| Vm | Early learning | 21.96 ± 0.60<br>(n = 19 neurons in 5 mice) | 19.00 ± 0.72<br>(n = 38 neurons in 10 mice) | 18.28 ± 0.74<br>(n = 38 neurons in 10 mice) | 22.29 ± 0.76<br>(n = 29 neurons in 10 mice) | 16.52 ± 0.80<br>(n = 29 neurons in 10 mice) | 15.44 ± 0.68<br>(n = 29 neurons in 10 mice) |
| | Late learning | 21.16 ± 0.55<br>(n = 17 neurons in 4 mice) | 16.95 ± 0.78<br>(n = 27 neurons in 8 mice) | 19.02 ± 1.14<br>(n = 27 neurons in 8 mice) | 22.29 ± 1.12<br>(n = 22 neurons in 8 mice) | 15.35 ± 0.78<br>(n = 22 neurons in 8 mice) | 14.17 ± 0.35<br>(n = 22 neurons in 8 mice) |
| FR | Early learning | 23.26 ± 0.49<br>(n = 19 neurons in 5 mice) | 18.44 ± 0.75<br>(n = 38 neurons in 10 mice) | 17.15 ± 0.72<br>(n = 38 neurons in 10 mice) | 21.89 ± 0.92<br>(n = 29 neurons in 10 mice) | 16.23 ± 0.49<br>(n = 29 neurons in 10 mice) | 15.10 ± 0.75<br>(n = 29 neurons in 10 mice) |
| | Late learning | 21.50 ± 0.99<br>(n = 17 neurons in 4 mice) | 16.30 ± 0.75<br>(n = 27 neurons in 8 mice) | 18.60 ± 1.30<br>(n = 27 neurons in 8 mice) | 21.09 ± 1.73<br>(n = 22 neurons in 8 mice) | 16.21 ± 1.00<br>(n = 22 neurons in 8 mice) | 14.65 ± 0.65<br>(n = 22 neurons in 8 mice) |

**Table S2.**
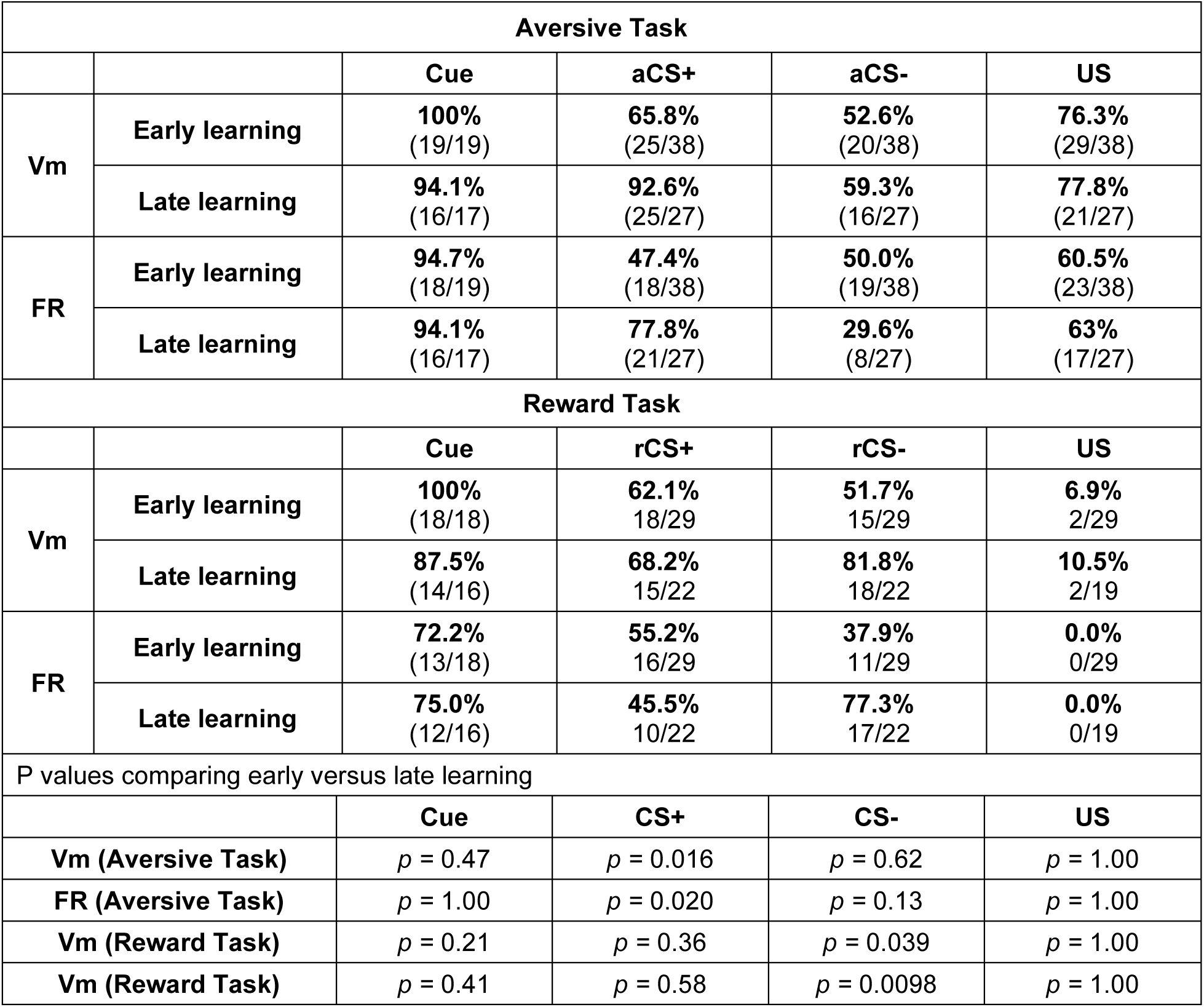
Summary of the fraction of neurons modulated during the immediate window by task-relevant events in the two tasks. Values indicate the number of modulated neurons over the total number of neurons tested. P values are comparisons with Fisher’s exact tests.

**Table S3.** Summary of the fraction of neurons discriminating two CSs in the two tasks. Values indicate the number of neurons showing significantly different responses between CS+ and CS− trials over the total number of neurons tested.

| **Aversive task (immediate window)** |  |  |  |  |
| --- | --- | --- | --- | --- |
|  | **Early learning** |  | **Late learning** |  |
|  | **Vm** | **FR** | **Vm** | **FR** |
| **aCS+ > aCS-** | **2.6%**<br>(1/38) | **7.9%**<br>(3/38) | **59.3%**<br>(16/27) | **51.9%**<br>(14/27) |
| **aCS+ < aCS-** | **5.3%**<br>(2/38) | **0.0%**<br>(0/38) | **0.0%**<br>(0/27) | **0.0%**<br>(0/27) |
| **Reward task (immediate window)** |  |  |  |  |
|  | **Early learning** |  | **Late learning** |  |
|  | **Vm** | **FR** | **Vm** | **FR** |
| **rCS+ > rCS-** | **6.9%**<br>(2/29) | **0.0%**<br>(0/29) | **0.0%**<br>(0/22) | **0.0%**<br>(0/22) |
| **rCS+ < rCS-** | **10.3%**<br>(3/29) | **3.5%**<br>(1/29) | **45.5%**<br>(10/22) | **18.2%**<br>(4/22) |
| **Aversive task (delayed window)** |  |  |  |  |
|  | **Early learning** |  | **Late learning** |  |
|  | **Vm** | **FR** | **Vm** | **FR** |
| **aCS+ > aCS-** | **7.9%**<br>(3/38) | **0.0%**<br>(0/38) | **3.7%**<br>(1/27) | **3.7%**<br>(1/27) |
| **aCS+ < aCS-** | **2.6%**<br>(1/38) | **0.0%**<br>(1/38) | **11.1%**<br>(3/27) | **3.7%**<br>(1/27) |
| **Reward task (delayed window)** |  |  |  |  |
|  | **Early learning** |  | **Late learning** |  |
|  | **Vm** | **FR** | **Vm** | **FR** |
| **rCS+ > rCS-** | **6.9%**<br>(2/29) | **3.5%**<br>(1/29) | **9.1%**<br>(2/22) | **9.1%**<br>(2/22) |
| **rCS+ < rCS-** | **0.0%**<br>(0/29) | **0.0%**<br>(0/29) | **4.6%**<br>(1/22) | **0.0%**<br>(0/22) |

**Table S4.**
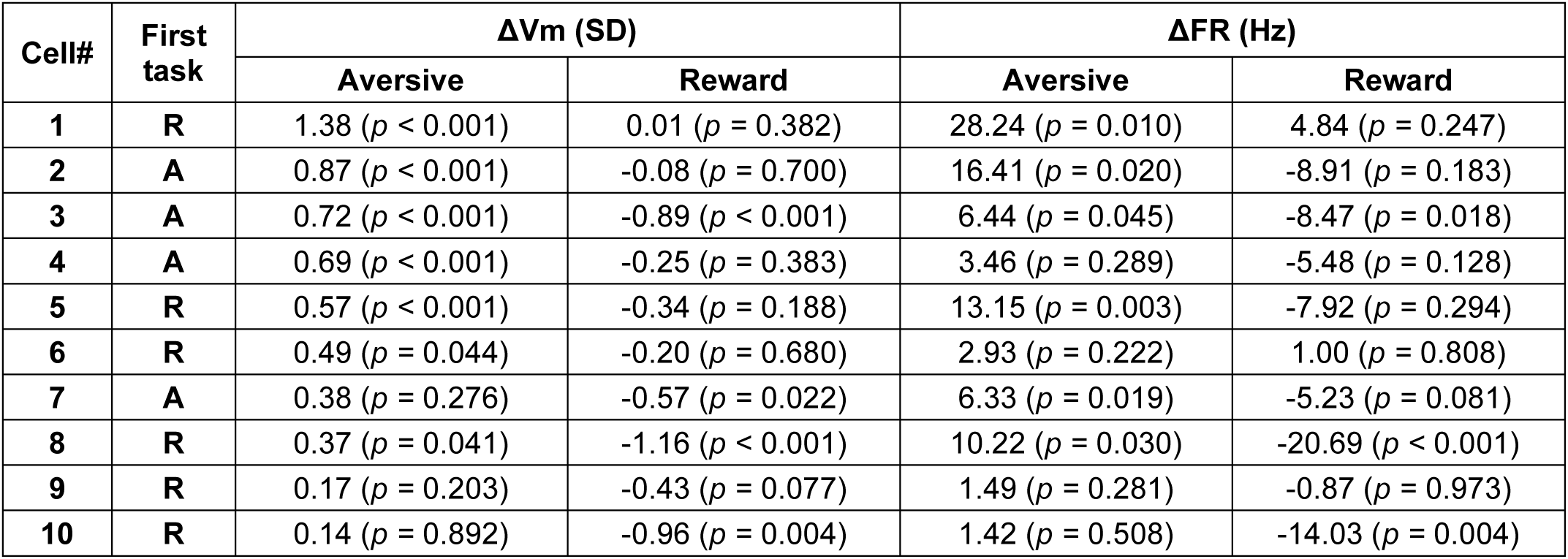
Individual neuron discrimination of CS+ and CS- across the 10 neurons recorded in both tasks. Values indicate the difference between the mean of immediate response in CS+ and CS- trials for each neuron (ΔVm and ΔFR, CS+ minus CS-). P values indicate trial-wise comparisons between CS+ and CS- trials (Mann-Whitney U tests). Cell ordering matches Fig. 5A.

| **Cell#** | **First task** | **$\Delta Vm$ (SD)** |  | **$\Delta FR$ (Hz)** |  |
| --- | --- | --- | --- | --- | --- |
|  |  | **Aversive** | **Reward** | **Aversive** | **Reward** |
| **1** | **R** | 1.38 ($p < 0.001$) | 0.01 ($p = 0.382$) | 28.24 ($p = 0.010$) | 4.84 ($p = 0.247$) |
| **2** | **A** | 0.87 ($p < 0.001$) | -0.08 ($p = 0.700$) | 16.41 ($p = 0.020$) | -8.91 ($p = 0.183$) |
| **3** | **A** | 0.72 ($p < 0.001$) | -0.89 ($p < 0.001$) | 6.44 ($p = 0.045$) | -8.47 ($p = 0.018$) |
| **4** | **A** | 0.69 ($p < 0.001$) | -0.25 ($p = 0.383$) | 3.46 ($p = 0.289$) | -5.48 ($p = 0.128$) |
| **5** | **R** | 0.57 ($p < 0.001$) | -0.34 ($p = 0.188$) | 13.15 ($p = 0.003$) | -7.92 ($p = 0.294$) |
| **6** | **R** | 0.49 ($p = 0.044$) | -0.20 ($p = 0.680$) | 2.93 ($p = 0.222$) | 1.00 ($p = 0.808$) |
| **7** | **A** | 0.38 ($p = 0.276$) | -0.57 ($p = 0.022$) | 6.33 ($p = 0.019$) | -5.23 ($p = 0.081$) |
| **8** | **R** | 0.37 ($p = 0.041$) | -1.16 ($p < 0.001$) | 10.22 ($p = 0.030$) | -20.69 ($p < 0.001$) |
| **9** | **R** | 0.17 ($p = 0.203$) | -0.43 ($p = 0.077$) | 1.49 ($p = 0.281$) | -0.87 ($p = 0.973$) |
| **10** | **R** | 0.14 ($p = 0.892$) | -0.96 ($p = 0.004$) | 1.42 ($p = 0.508$) | -14.03 ($p = 0.004$) |

**Table S5.** Statistical table for Fig. 3.

| Aversive task |  |  |  |  |  |  |
| --- | --- | --- | --- | --- | --- | --- |
| P values comparing evoked responses to pre-event baseline across neurons (Wilcoxon signed-rank tests). |  |  |  |  |  |  |
| Panel |  |  | Cue | aCS+ | aCS- | US |
| Fi | Vm (immediate) | Early learning | $p < 0.001$ | $p < 0.001$ | $p < 0.001$ | $p < 0.001$ |
| | | Late learning | $p < 0.001$ | $p < 0.001$ | $p < 0.001$ | $p < 0.001$ |
| Fii | FR (immediate) | Early learning | $p < 0.001$ | $p < 0.001$ | $p < 0.001$ | $p < 0.001$ |
| | | Late learning | $p < 0.001$ | $p < 0.001$ | $p < 0.001$ | $p < 0.001$ |
| Gi | Vm (delayed) | Early learning | $p = 0.040$ | $p = 0.94$ | $p = 0.61$ | $p = 0.005$ |
| | | Late learning | $p = 0.002$ | $p = 0.005$ | $p = 0.25$ | $p = 0.50$ |
| Gii | FR (delayed) | Early learning | $p = 0.11$ | $p = 0.52$ | $p = 0.74$ | $p = 0.001$ |
| | | Late learning | $p = 0.004$ | $p = 0.99$ | $p = 0.15$ | $p = 0.44$ |
| P values comparing early versus late learning across neurons (Mann-Whitney U tests). |  |  |  |  |  |  |
| Panel |  |  | Cue | aCS+ | aCS- | US |
| Fi | Vm (immediate) | | $p = 0.011$ | $p < 0.001$ | $p = 0.84$ | $p = 0.48$ |
| Fii | FR (immediate) | | $p = 0.014$ | $p < 0.001$ | $p = 0.95$ | $p = 1.00$ |
| Gi | Vm (delayed) | | $p = 0.04$ | $p = 0.03$ | $p = 0.59$ | $p = 0.09$ |
| Gii | FR (delayed) | | $p = 0.25$ | $p = 0.65$ | $p = 0.53$ | $p = 0.06$ |

| Reward task |  |  |  |  |  |  |
| --- | --- | --- | --- | --- | --- | --- |
| P values comparing evoked responses to pre-event baseline across neurons (Wilcoxon signed-rank tests). |  |  |  |  |  |  |
| Panel |  |  | Cue | rCS+ | rCS- | US |
| Mi | Vm (immediate) | Early learning | $p < 0.001$ | $p < 0.001$ | $p < 0.001$ | $p = 0.42$ |
| | | Late learning | $p < 0.001$ | $p < 0.001$ | $p < 0.001$ | $p = 0.37$ |
| Mii | FR (immediate) | Early learning | $p < 0.001$ | $p < 0.001$ | $p = 0.001$ | $p = 0.47$ |
| | | Late learning | $p < 0.001$ | $p < 0.001$ | $p < 0.001$ | $p = 0.46$ |
| Ni | Vm (delayed) | Early learning | $p = 0.61$ | $p = 0.26$ | $p = 0.94$ | $p = 0.004$ |
| | | Late learning | $p = 0.028$ | $p = 0.042$ | $p = 0.49$ | $p = 0.08$ |
| Nii | FR (delayed) | Early learning | $p = 0.60$ | $p = 0.42$ | $p = 0.63$ | $p = 0.15$ |
| | | Late learning | $p = 0.51$ | $p = 0.07$ | $p = 0.45$ | $p = 0.25$ |
| P values of comparing early versus late learning across neurons (Mann-Whitney U tests). |  |  |  |  |  |  |
| Panel |  |  | Cue | rCS+ | rCS- | US |
| Mi | Vm (immediate) | | $p = 0.034$ | $p = 0.65$ | $p < 0.001$ | $p = 0.62$ |
| Mii | FR (immediate) | | $p = 0.024$ | $p = 0.54$ | $p < 0.001$ | $p = 0.45$ |
| Ni | Vm (delayed) | | $p = 0.021$ | $p = 0.010$ | $p = 0.32$ | $p = 0.87$ |
| Nii | FR (delayed) | | $p = 0.78$ | $p = 0.13$ | $p = 0.49$ | $p = 0.11$ |

**Table S6.**
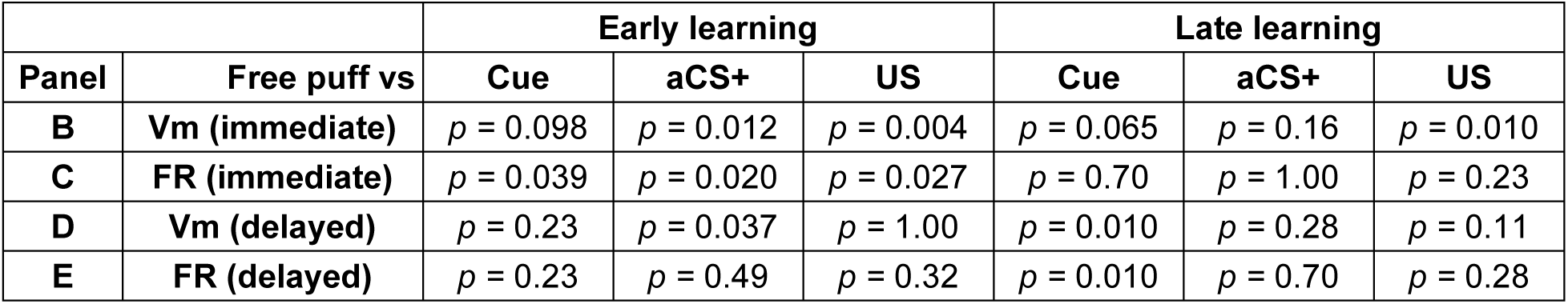
Statistical Table for Fig. S4. P values of comparing the evoked responses by free-puff versus other events across neurons (Wilcoxon signed-rank tests).

| Panel | Free puff vs | Early learning |  |  | Late learning |  |  |
| --- | --- | --- | --- | --- | --- | --- | --- |
|  |  | Cue | aCS+ | US | Cue | aCS+ | US |
| B | Vm (immediate) | $p = 0.098$ | $p = 0.012$ | $p = 0.004$ | $p = 0.065$ | $p = 0.16$ | $p = 0.010$ |
| C | FR (immediate) | $p = 0.039$ | $p = 0.020$ | $p = 0.027$ | $p = 0.70$ | $p = 1.00$ | $p = 0.23$ |
| D | Vm (delayed) | $p = 0.23$ | $p = 0.037$ | $p = 1.00$ | $p = 0.010$ | $p = 0.28$ | $p = 0.11$ |
| E | FR (delayed) | $p = 0.23$ | $p = 0.49$ | $p = 0.32$ | $p = 0.010$ | $p = 0.70$ | $p = 0.28$ |

**Table S7.** Statistical Table for Fig. S6. P values of comparing the evoked responses by free-reward versus other events across neurons (Wilcoxon signed-rank tests).

| Panel | Free reward vs | Early learning |  |  | Late learning |  |  |
| --- | --- | --- | --- | --- | --- | --- | --- |
|  |  | Cue | rCS+ | US | Cue | rCS+ | US |
| B | Vm (immediate) | $p = 0.36$ | $p = 0.76$ | $p < 0.001$ | $p = 0.74$ | $p = 0.89$ | $p < 0.001$ |
| C | FR (immediate) | $p = 0.95$ | $p = 0.30$ | $p = 0.003$ | $p = 0.95$ | $p = 0.46$ | $p = 0.003$ |
| D | Vm (delayed) | $p = 0.035$ | $p = 0.14$ | $p = 1.00$ | $p = 0.048$ | $p = 0.033$ | $p = 0.50$ |
| E | FR (delayed) | $p = 0.24$ | $p = 0.030$ | $p = 0.17$ | $p = 0.24$ | $p = 0.50$ | $p = 0.79$ |

**Table S8.**
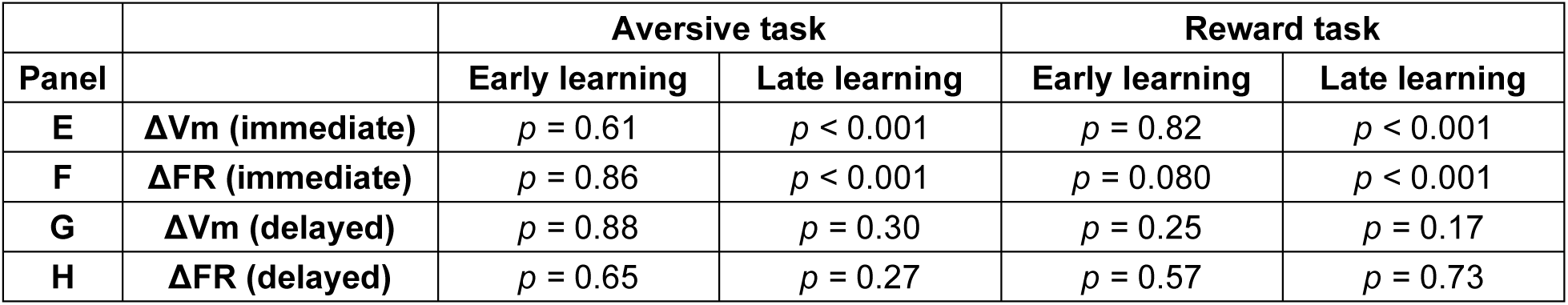
Statistical table for Fig. 4E-H. P values comparing ΔVm (CS+ minus CS-) and ΔFR against zero across neurons (Wilcoxon signed-rank tests).

**Table S9.** Statistical table for Fig. S7. P values comparing responses to evoked by CS+ and CS- across neurons (Wilcoxon signed-rank tests) in both tasks.

|  | Panel | Immediate window |  | Panel | Delayed window |  |
| --- | --- | --- | --- | --- | --- | --- |
|  |  | Aversive | Reward |  | Aversive | Reward |
| Vm (early learning) | Bi | $p = 0.61$ | $p = 0.84$ | Bii | $p = 0.88$ | $p = 0.25$ |
| Vm (late learning) | Di | $p < 0.001$ | $p < 0.001$ | Dii | $p = 0.67$ | $p = 0.61$ |
| FR (early learning) | Fi | $p = 0.86$ | $p = 0.15$ | Fii | $p = 0.87$ | $p = 0.57$ |
| FR (late learning) | Hi | $p < 0.001$ | $p = 0.006$ | Hii | $p = 0.27$ | $p = 0.34$ |

**Table S10.** Statistical table for Fig. 4I-L. P values comparing trial-start-cue-evoked responses to pre-onset baseline across neurons (Wilcoxon signed-rank tests), and trial-start-cue-evoked responses between aversive and reward tasks (Mann-Whitney U tests).

| Panel |  | Early learning |  |  | Late learning |  |  |
| --- | --- | --- | --- | --- | --- | --- | --- |
|  |  | Aversive | Reward | Aversive vs Reward | Aversive | Reward | Aversive vs Reward |
| I | Vm (immediate) | $p < 0.001$ | $p < 0.001$ | $p = 0.13$ | $p < 0.001$ | $p < 0.001$ | $p = 0.043$ |
| J | FR (immediate) | $p < 0.001$ | $p < 0.001$ | $p = 0.43$ | $p < 0.001$ | $p < 0.001$ | $p = 0.048$ |
| K | Vm (delayed) | $p = 0.11$ | $p = 0.84$ | $p = 0.14$ | $p = 0.006$ | $p = 0.070$ | $p = 0.19$ |
| L | FR (delayed) | $p = 0.007$ | $p = 0.49$ | $p = 0.06$ | $p = 0.001$ | $p = 0.796$ | $p = 0.004$ |

**Table S11.** Statistical Table for Fig. 5.

P values comparing between various CS pairs across neurons (Wilcoxon signed-rank tests).
|  |  | aCS+ vs |  |  | aCS- vs |  | rCS+ vs |
| --- | --- | --- | --- | --- | --- | --- | --- |
| Panel |  | aCS- | rCS+ | rCS- | rCS+ | rCS- | rCS- |
| J | Vm (immediate) | $p = 0.002$ | $p = 0.037$ | $p = 0.85$ | $p = 0.43$ | $p = 0.013$ | $p = 0.003$ |
| K | FR (immediate) | $p = 0.002$ | $p = 0.027$ | $p = 0.70$ | $p = 0.92$ | $p = 0.003$ | $p = 0.013$ |
| L | Vm (delayed) | $p = 0.77$ | $p = 0.49$ | $p = 0.70$ | $p = 0.43$ | $p = 0.56$ | $p = 0.85$ |
| M | FR (delayed) | $p = 0.70$ | $p = 0.38$ | $p = 0.92$ | $p = 0.43$ | $p = 1.00$ | $p = 0.28$ |

P values comparing trial-start-cue-evoked response across tasks (Wilcoxon signed-rank tests).

| Panel |  | Cue in aversive vs Cue in reward |
| --- | --- | --- |
| O | Vm (immediate) | $p = 0.027$ |
| P | FR (immediate) | $p = 0.13$ |
| Q | Vm (delayed) | $p = 0.56$ |
| R | FR (delayed) | $p = 0.002$ |

**Table S12.**
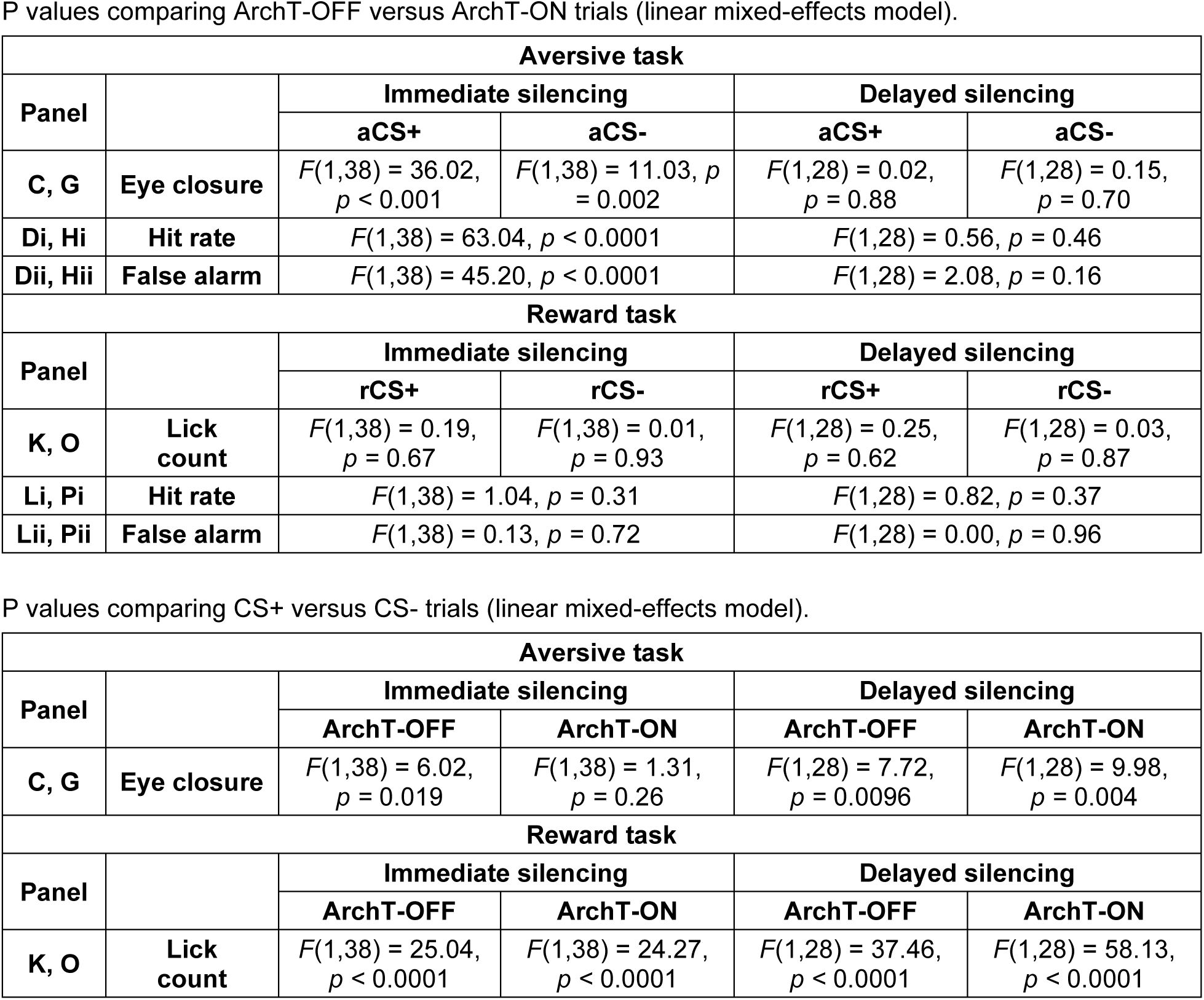
Statistical Table for Fig. 6.

